# MetaMiner: Streamlined GUI Tool for Retrieving, Normalizing and Exploring Metadata

**DOI:** 10.1101/2025.08.20.666107

**Authors:** Jaykumar Kiritkumar Patel, Ravikrishnan Elangovan

## Abstract

**Summary:** Over the past decade, publicly available genomic data has expanded exponentially. However, the usability of these genomes heavily depends on their associated metadata. Metadata provides context about the data that from where, when and how the data was obtained. When not consistent, it limits the re-usage of the available public data and prevents data’s integration into broader genomic analyses. Here, we introduce MetaMiner, a GUI tool designed to retrieve, normalize, and interactively visualize metadata associated with prokaryotic genomes. MetaMiner streamlines the normalization of key metadata elements such as geographical locations, isolation sources, and sequencing technologies. To assess its usability, we used metadata of *Acinetobacter baumannii* genomes and compared it with NCBI’s data processing tools and manually curated datasets. MetaMiner demonstrated superior performance in data transformation and showed high classification accuracy, with only minor discrepancies arising from recent geopolitical changes or ambiguous submissions. This cleaned and standardized metadata is integrated into an interactive dashboard, which allows users to explore data holistically through visualizations and custom filtering options. The dashboard has interactive plots related to submission date trend, geographical origins, sequencing methods, genome quality scores (e.g., ANI > 95, BUSCO > 95), coverage, etc. Overall, MetaMiner minimizes manual curation efforts of genomic metadata and enhances the accessibility, and re-usability of publicly available genomes.

**Availability and Implementation:** The executables and source code are both freely available to use at https://github.com/prekijpatel/MetaMiner for non-commercial use. Detailed documentation is available at https://github.com/prekijpatel/MetaMiner/wiki.

## Background

High-throughput sequencing technologies has revolutionised various fields of biology, leading to exponential increase in omics data generation (Reuter *et al*., 2015). Resultingly, genome submissions to public databases, such as GenBank and the European Nucleotide Archive (ENA), have increased significantly in recent years, highlighting the rapid rise of sequencing projects worldwide (Sayers *et al*., 2025; O’Cathail *et al*., 2025). The value of this vast genomic datasets depends not only on the sequences themselves but also on the quality and completeness of the accompanying metadata. Metadata—information about the conditions and context in which the data was collected—is extremely helpful in designing new hypotheses and experiments by re-using old data (Caliskan *et al*., 2023). This is especially important in fields like comparative genomics. Conducting comparative genomic studies and inferring from them often depends on well-organized and standardised metadata (Bornstein *et al*., 2023; Moustafa *et al*., 2020).

Due to the adoption of FAIR (Findable, Accessible, Interoperable, and Reusable) principles (Wilkinson *et al*., 2016; Musen *et al*., 2022) and recent developments, metadata retrieval from public repositories has become easier. The retrieval is usually through command-line tools or FTP sites (O’Leary *et al*., 2024; Gálvez-Merchán *et al*., 2023). Despite these advancements in availability of metadata, the quality of metadata still remains a limiting factor in its applications. To address this, the Genomic Standards Consortium (GSC) has established standards such as MIGS (Minimum Information about a Genome Sequence), MIMS (Minimum Information about a Metagenome Sequence), MISAG (Minimum Information about a Single Amplified Genome), and MIMAG (Minimum Information about a Metagenome-Assembled Genome) (Field *et al*., 2008; Bowers *et al*., 2017). While these frameworks offer valuable guidance for metadata standardisation, many fields still allow freeform text entries without validation. This often leads to unintentional errors, such as misspellings, missing entries, inconsistency, and over-explanation, which diminishes the usability and reliability of the data. These errors are even more common when multiple researchers and/or studies are involved. In addition to this, genomic data’s high dimensionality makes it even more complex as each dataset, varying from human-associated pathogens to environmental symbiont, may have its own unique metadata needs and accompanying challenges–which makes the standardisation/normalisation of metadata even more perplexing.

Improving metadata standardisation and quality has the potential to advance genomics research to a great extent. High-quality metadata allows researchers to combine data from different labs and studies, facilitating collaborative research and enabling experiments that are beyond the scope of individual labs (Ryan *et al*., 2021; Rajesh *et al*., 2021). Standardisation of metadata also makes it machine-actionable, which enables automated processing and further simplification of metadata without manual labour (Batista *et al*., 2022). Overall, standardised metadata immensely aids in efficient data discovery, reuse, and interpretation.

There are resources which are widely recognised for providing uniformly analysed genomes and/or curated metadata, such as EnteroBase (Dyer *et al*., 2024), the Genomes OnLine Database (Mukherjee *et al*., 2025), the National Microbiome Data Collaborative (NMDC) (National Microbiome Data Collaborative), and BakRep (Fenske *et al*., 2024). However, as these repositories often focus on distinct taxa or specific research goals, their metadata may have limitations in general usability for broader comparative studies.

To address these gaps, we introduce MetaMiner, a GUI-based tool designed to streamline metadata retrieval, cleaning, and standardisation for prokaryotic genomes from NCBI datasets. It systematically normalizes key metadata elements such as geographical location, isolation source, sequencing technology, and other assembly parameters. Additionally, it identifies and removes redundancies within datasets, and outputs a cleaner, programmatically curated metadata set, which, with minimal cross-checking, results in reliable and standardised metadata. Beyond metadata cleaning, MetaMiner features an interactive visualisation dashboard powered by Dash, allowing users to dynamically explore, filter, and analyse metadata in an intuitive interface.

## Methodology

### Metadata Retrieval and Processing

The *Acinetobacter baumannii* metadata was obtained on February 3, 2025. The selection of *Acinetobacter baumannii* was random and serves as a representative example. The command-line downloaded metadata was downloaded in JSON-lines format using NCBI’s *datasets*.*exe (v16*.*40*.*1)*. The *dataformat*.*exe (v16*.*40*.*1)* from NCBI was used to format the JSON-line from a webpage, JSON-line from command-line tool, and JSON from MetaMiner. This was then compared to MetaMiner formatted versions. The log for metadata retrieval and formatting is available in *Supplementary file 1*. The time estimation for loading formatted TSV files was calculated using the *timeit* module for five iterations.

### Comparing Metadata Normalization

MetaMiner normalised data was compared with the manually/peer-curated normalisations. The raw metadata that was normalised was same as the one retrieved using MetaMiner on 3^rd^ February 2025. The code used for comparison and plotting the results is available in *Supplementary file 2*.

## Results

### Design

MetaMiner is a Python-based tool developed for the retrieval, standardisation, and visualisation of prokaryotic metadata. It makes use of various libraries and custom-made databases to process metadata efficiently. The tool is designed to handle both scenarios where pre-downloaded metadata is already available with the user and another where the user needs to download the needed metadata from the NCBI. The metadata retrieval from NCBI is enabled by the datasets command-line tool. MetaMiner transforms the raw JSON/JSONL to a structured, tabular format, making it easier to work with in large-scale analyses. Once the metadata is transformed, preprocessing steps are applied to remove redundant entries from the formatted metadata, specifically, for assemblies that include both RefSeq and GenBank annotations, only the metadata from RefSeq assembly is retained for subsequent processing while retaining the raw metadata in a separate file.

To standardise geographical location data, MetaMiner utilises a combination of text-based normalisation and geospatial tools such as *PyCountry* and *Geopy’s Nominatim*. The tool extracts and maps geographical information to country- and state-levels based on raw metadata entries. It is important to note that MetaMiner does not perform sub-state level normalisation; therefore, even if district-level information is available, it is aggregated at the state level to ensure uniformity.

Normalisation of isolation source metadata is done by utilising rapidfuzz library on certain keywords and localised reference database. The algorithm leverages three metadata fields, `*host*`, `*host_disease*`, and *`isolation_source`*, to categorize the isolation sources. Each raw metadata entry, defined as a combination of the `*host*`, `*host_disease*`, and `*isolation_source*` fields, is matched against an extensive internal dictionary. This dictionary comprises standardized terms, synonyms, common misspellings, unusual abbreviations, and other irregular forms frequently encountered in raw datasets. Matching is conducted using *rapidfuzz*, which applies fuzzy string-matching techniques guided by empirically determined thresholds to account for near matches and minor variations. Based on the input, the algorithm categorizes isolation source into a four-level classification. The first level broadly categorizes the entry by host type such as, Hospital, Animal, Environmental, Laboratory, among others. The remaining three levels of classification are host-specific and vary depending on the assigned host category. The detailed overview of levelled classification categories and algorithm is illustrated in Figure 1 and 2, respectively.

**Figure 1.**
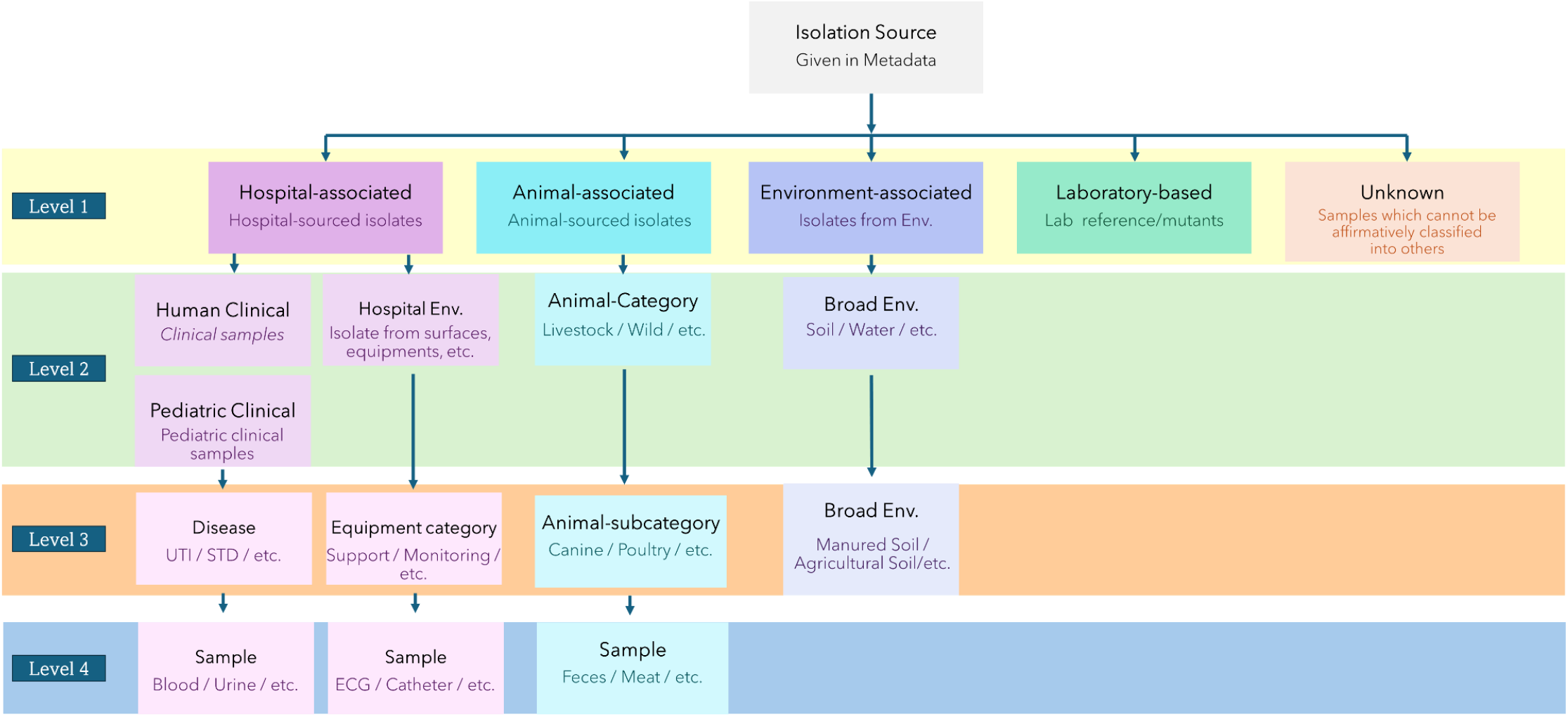
Classification scheme illustrating the hierarchical levels in the normalized isolation source data. *The four levels are depicted using distinct colors. In case of Hospital-associated isolates, unlike others, the level 2 has 3 separate and independent categories which are classified differently in proceeding levels*.

**Figure 2.**
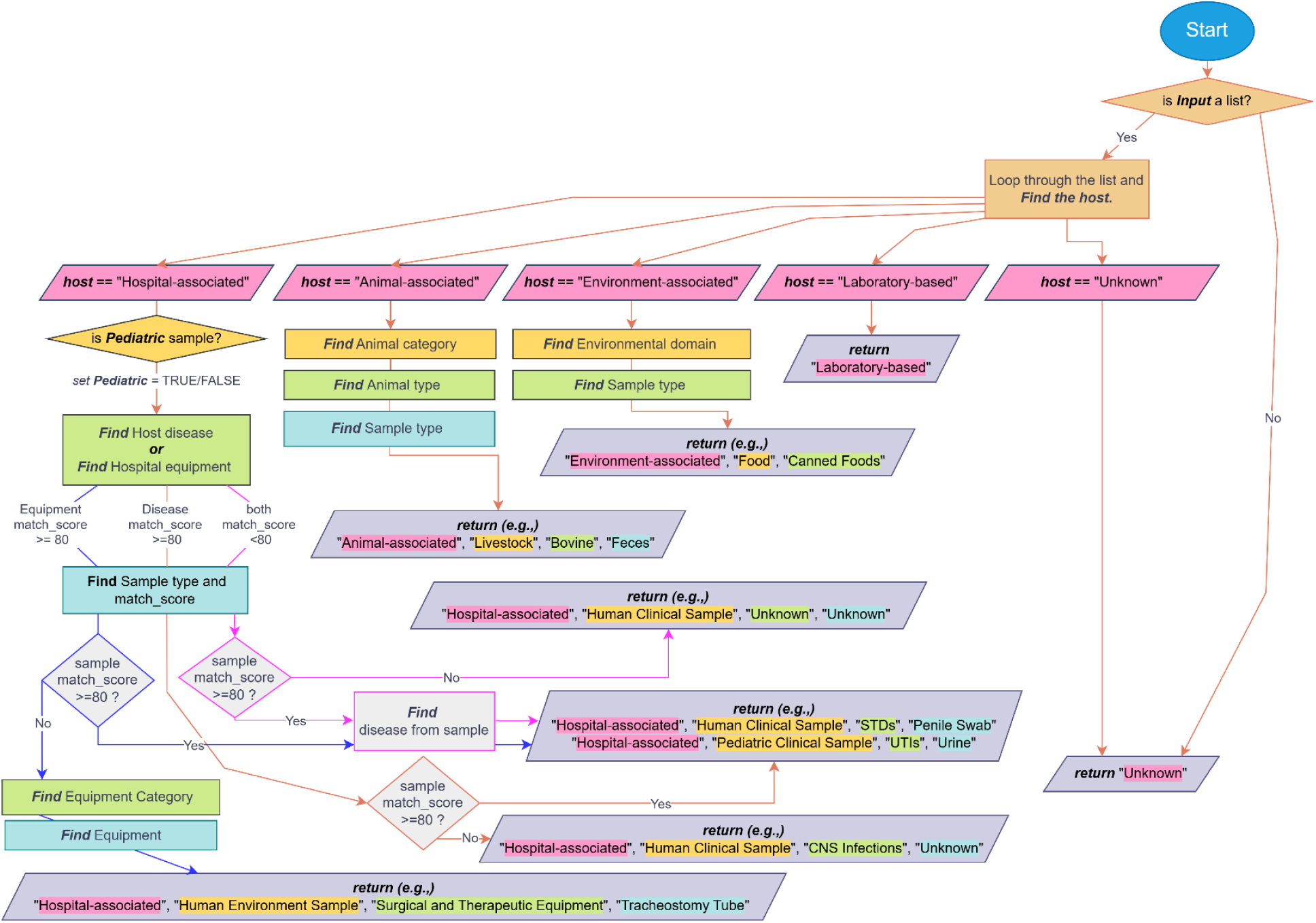
Flowchart depicting the algorithm for normalizing isolation source metadata. The different level of information is retrieved at various steps of the algorithm. The examples of final retrieved results are shown in purple parallelograms. The text-background colour corresponds to the process step (box color) from which the information was obtained.

Sequencing technology used to generate assembly data is classified by identifying key terms such as Illumina, Nanopore, DNBseq, MiSeq, etc. Based on them, MetaMiner categorises sequencing technologies into various short-read, long-read, or hybrid sequencing platforms. The representative examples of these normalizations are depicted in Table 1. In addition to this, other assembly-related parameters, such as gene counts, ANI, BUSCO, coverage, etc., are also re-formatted. All of this is streamed into an in-built dashboard which allows user to interact with metadata visually, providing functionalities such as filtering specific subsets of data, exploring trends through dynamic plots, and saving customized views or filtered datasets for downstream analysis and reporting purposes.

**Table 1.** Examples of Normalization performed by MetaMiner. The table illustrates the transformation of heterogeneous metadata into standard formats across three metadata fields: Geographical locations, Isolation source and Sequencing Technology.

| Geographical Location |  |  |  |  |
| --- | --- | --- | --- | --- |
| Raw data | Normalized Data |  |  |  |
| Raw *geo_loc* data | Identified Country | Country Code | Identified State | State ISO Code |
| India: Pune; BJ Medical College | India | IND | Maharashtra | IN-MH |
| Brasil: São Paulo | Brazil | BRA | São Paulo | BR-SP |
| USA: CA | United States | USA | California | US-CA |
| Germany: Bayern | Germany | DEU | Bavaria | DE-BY |
| Wuhan Institute of Virology | China | CHN | Hubei Sheng | CN-HB |
| Isolation Source |  |  |  |  |
| Raw data | Normalized Data |  |  |  |
| Raw *[host, host_disease, isolation_source]* data | Identified host | Identified Source Category | Identified Source | Sample |
| [Homo sapiens, UTIs, Urine Sample] | Hospital-associated | Human Clinical Sample | UTIs | Urine |
| [Infant, Severe Sepsis case, Blood] | Hospital-associated | Pediatric Clinical Sample | Sepsis/Septic Shock | Blood |
| [, , Ventilator tube tip] | Hospital-associated | Hospital Environment Sample | Surgical and Therapeutic Equipment | Ventilator |
| [Dog, ,wound swab] | Animal-associated | Companion | Canine | Swab |
| [, ,Chicken Meatball] | Animal-associated | Livestock | Poultry | Meat/Organ |
| [, ,Bootswab from poultry] | Animal-associated | Livestock | Poultry | Breeding/Hospital Environment Sample |
| [Synthetic variant of Salmonella Typhi] | Laboratory-based |  |  |  |
| [, ,Fecal Sample] | Unknown |  |  |  |
| [, ,Municipal wastewater sludge] | Environment-associated | Water | Wastewater sludge |  |
| Sequencing Technology |  |  |  |  |
| Raw data | Normalized Data |  |  |  |
| Raw *Sequencing Technology* Data | Standardized Sequencing Technology |  |  |  |
| miseq | Illumina |  |  |  |
| MiSeq (Illumina) | Illumina |  |  |  |
| MinION/GridION | Oxford Nanopore |  |  |  |
| Pacific Biosciences | PacBio |  |  |  |
| RSII | PacBio |  |  |  |
| myseq; minion | Illumina and Oxford Nanopore |  |  |  |
| nanopore + pacbio | Oxford Nanopore and PacBio |  |  |  |

### Validation of Metadata retrieval and processing

In order to test the efficiency of MetaMiner to retrieve and transform the data, raw metadata was retrieved for *Acinetobacter baumannii* assemblies using three different methods: (1) from the NCBI Datasets webpage, (2) via the NCBI Datasets command-line tool, and (3) using MetaMiner. Despite the varying retrieval methods and negligible differences in file size, the number of records in each dataset was identical (*Figure 3a*). This indicates that the data retrieved using MetaMiner is equivalent to that from NCBI, which corroborates with MetaMiner’s use of NCBI Datasets in the background.

**Figure 3.**
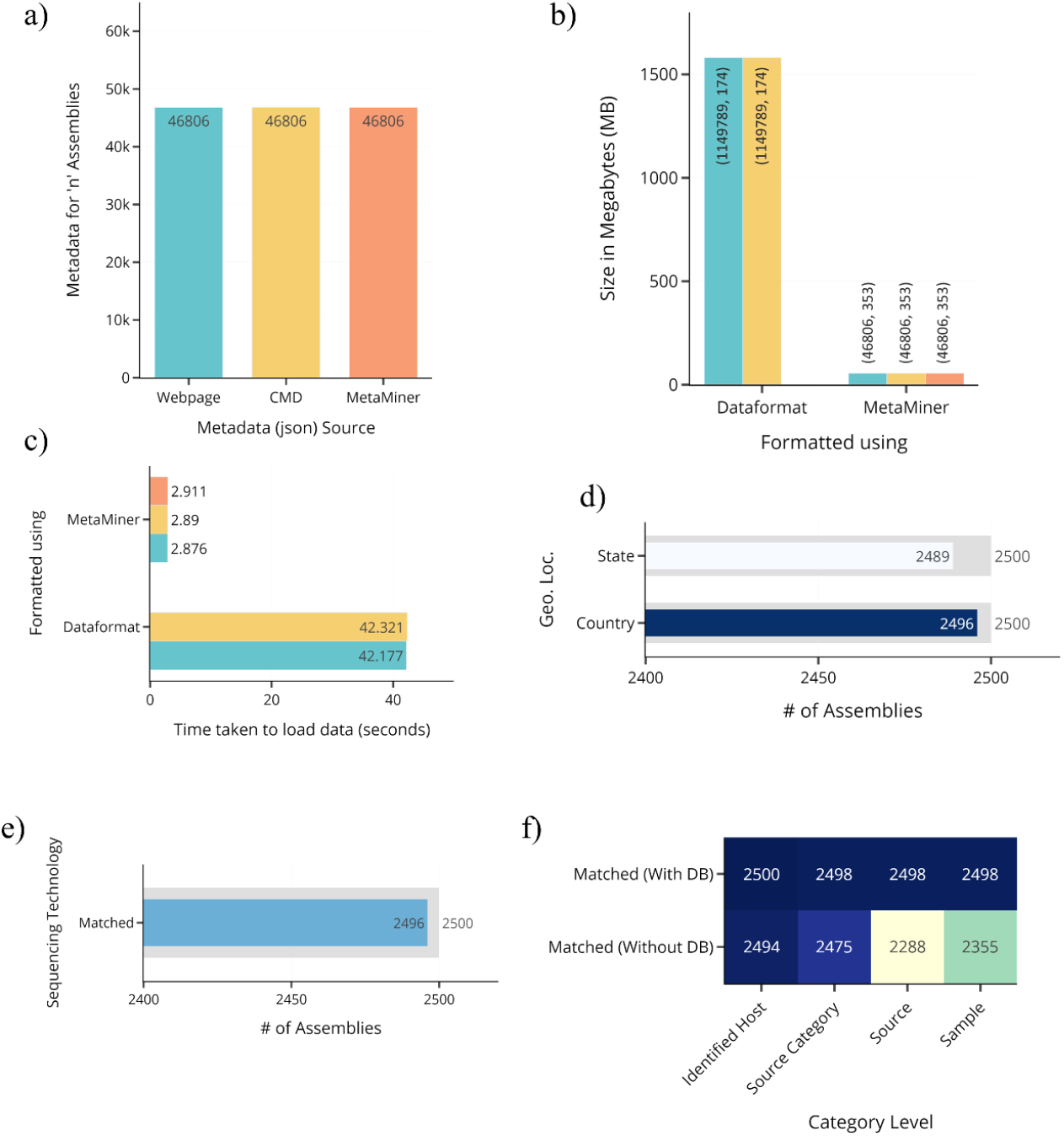
a) Number of assemblies for which metadata is available in json/jsonl file downloaded using various modalities. b) Size of the transformed metadata by NCBI dataformat and MetaMiner. The text near the bar represent the shape (row, column) of the transformed metadata table. c)Time taken to load the transformed metadata for further processing. d) Number of assemblies’ normalized geographical locations that matched with manually curated data. The manually curated data bar is shown in grey in back of each bar. e) Number of assemblies’ normalized sequencing technology data that matched with manually curated data. f) Number of assemblies’ categorized isolation source data that matched with manually curated data. The normalization was performed with and without the supporting database.

Next, to compare how MetaMiner processes raw metadata in JSON/JSONL format, we transformed the retrieved data into tab-separated values using both NCBI’s Dataformat and MetaMiner. The data formatted by MetaMiner is considerably smaller than the one formatted by NCBI Dataformat (*Figure 3b*). This difference arises from the distinct structuring methods; Dataformat creates a new row for each BioSample attribute, while MetaMiner consolidates all attributes into a single row. This approach significantly enhances the loading and processing times of the transformed data for further use (*Figure 3c*)..

Additionally, unlike Dataformat, MetaMiner can handle both JSON and JSON-lines formats. While NCBI Datasets and Dataformat offer many other features which MetaMiner does not, the advantages of MetaMiner in terms of transforming metadata are evident.

### Evaluation of Metadata Normalisation

A total of 2,500 assemblies were randomly selected from the retrieved and processed *A. baumannii* metadata. For these assemblies, data related to Geographical Location, Isolation Source, and Sequencing Technology were normalized using both manual curation and MetaMiner.

Geographical location normalization categorized the ‘geo_loc’ data from the raw data into Country and State information. MetaMiner demonstrated performance comparable to manual curation, correctly identifying 2,496 countries out of 2,500 assemblies (*Figure 3d*). The minor discrepancies stemmed from recent geopolitical changes (e.g., separation of certain countries) and ambiguous entries submitted by submitters, such as “Korea”, which could not be resolved by MetaMiner as North or South.

Like geographical locations, the normalization of sequencing technology used for genome sequencing also showed high level of accuracy (*Figure 3e*). The only exception was its inability to identify a single new sequencing technology, GenoLab M.

In terms of isolation source normalization, due to the high variability of isolation source entries, a curated database is used by default to support the categorization. This database is manually compiled from existing metadata of similar genomes and provides a reference set for normalization. However, to evaluate MetaMiner’s performance objectively, normalization was tested with and without the curated database. The without-database scenario simulates how the categorization would be handled when entirely new entries are encountered. *Figure 3f* presents the performance data, highlighting that MetaMiner successfully categorized most entries with the database across all levels. However, it encountered minor challenges in classifying ‘Source’ and ‘Sample’ categories in the absence of database support. Notably, even when MetaMiner is not able to categorise ‘Source’ and ‘Sample’ correctly, it still correctly identified the ‘host’ and ‘source category’ from which the sample was collected, demonstrating its robustness in metadata normalization at those levels if not more.

### Dash-based Dashboard

Once the data is cleaned, normalised, and standardised, it is processed to generate an interactive dashboard (Figure 4). This dashboard serves as the central interface for exploring the normalized dataset, offering both visual summaries and filtering capabilities. For example, normalized geographical locations are plotted into choropleth map (Figure 5), allowing users to identify regional sampling densities and trends. Sequencing platforms associated with each assembly are visualized through an interactive scatter plot (Figure 6), which along with coverage bar plots could be helpful in selecting highly accurate assemblies. To assess genome assembly quality, a scatter plot comparing L50 and N50 values is provided (Figure 7), helping to distinguish between high- and low-contiguity assemblies. The diversity of isolation sources is captured in a hierarchical tree map (Figure 8), where genomes are organized based on host association, isolation category, and sample type, offering an intuitive overview of the ecological and clinical contexts of the dataset.

**Figure 4.**
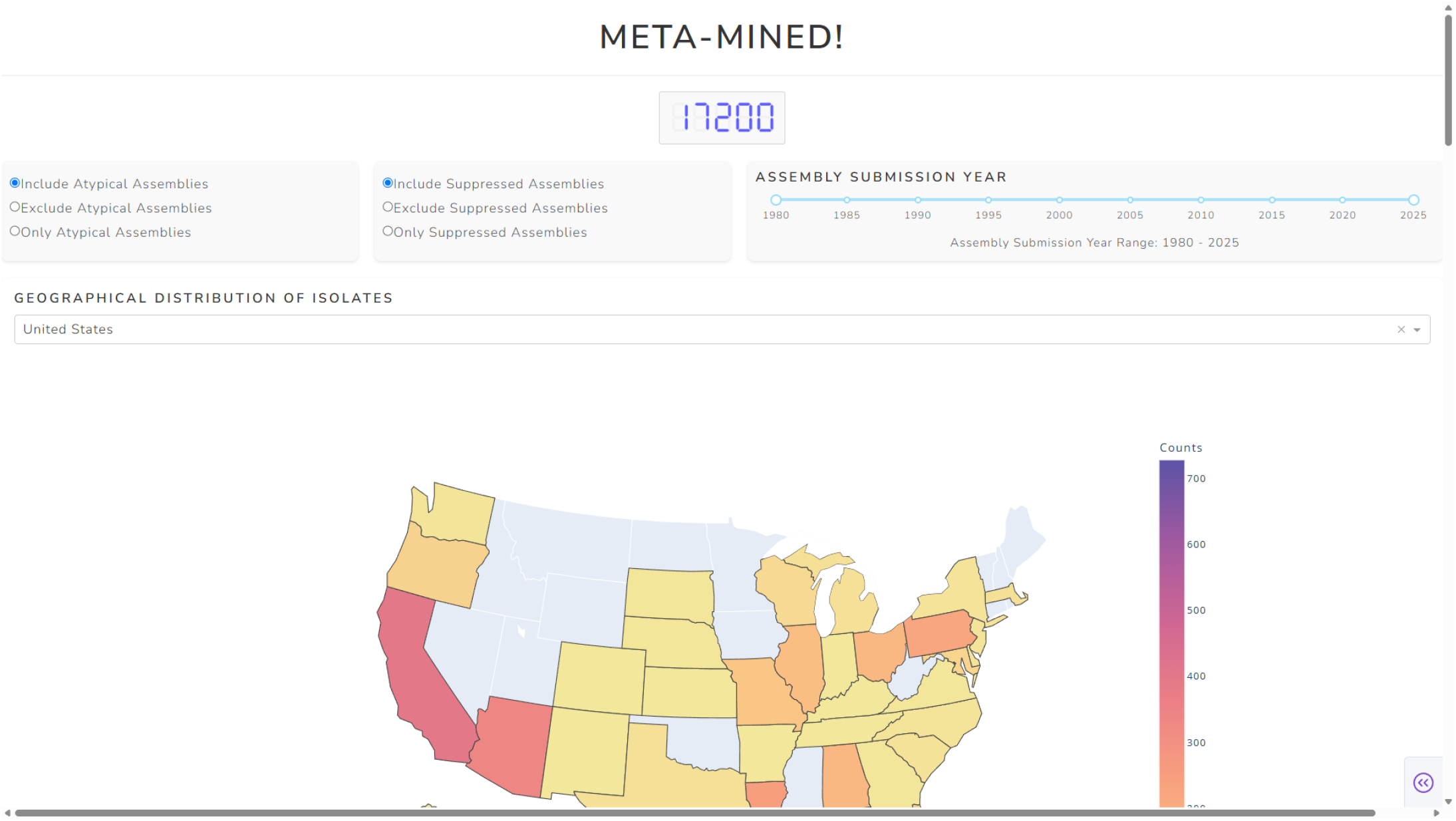
Top section of the Meta-Mined dashboard interface. *The screenshot displays the upper portion of the dashboard, including features such as LED showing total assembly number, radio buttons for including/excluding atypical and suppressed assemblies, and geographical distribution of isolates. Users can apply these filters along with other additional (not seen in the screenshot) ones—such as, sequencing technology, gene counts, isolation source, and genomic quality metrics, etc*.*—to explore and refine the dataset interactively*.

**Figure 5.**
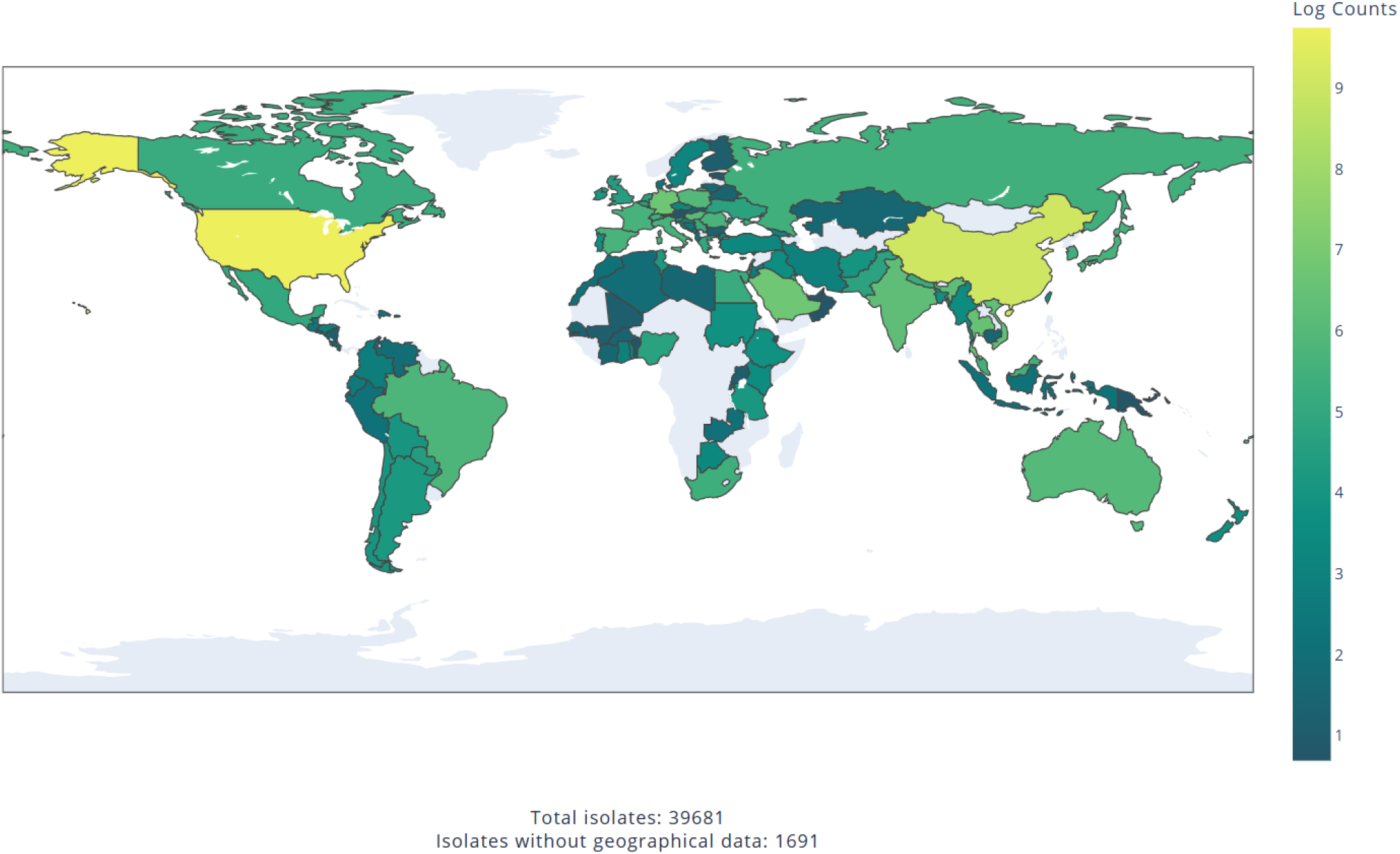
Choropleth map showing geographical distribution of assembles in Meta-Mined dashboard. *The static image represents the interactive choropleth map, which displays number of assemblies per country. In the bottom, it shows the number of total assemblies and number of assemblies for which geographical data is not available. In live dashboard, choropleth map is combined with a drop-down menu to select assemblies from specific country*.

**Figure 6.**
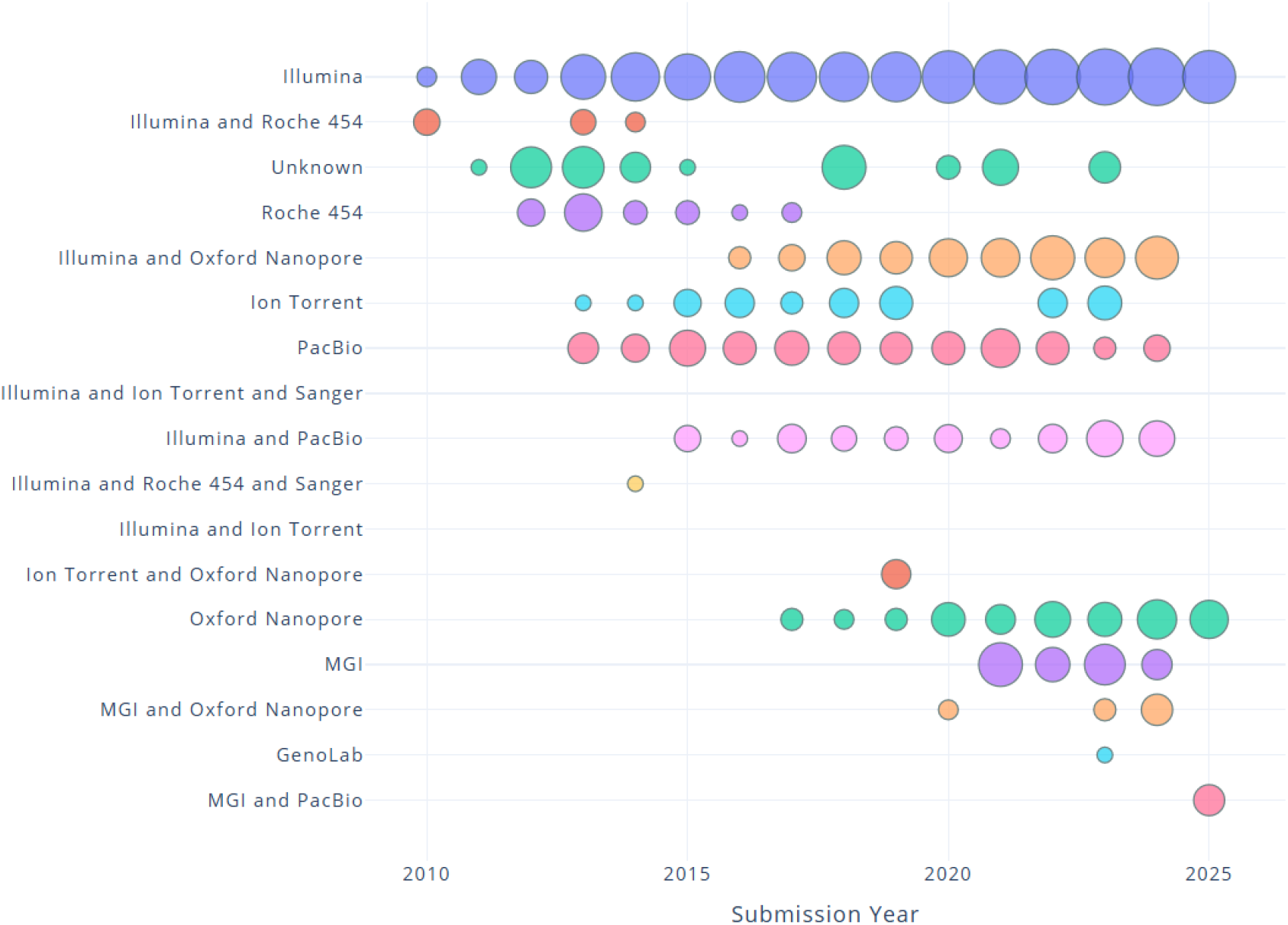
Scatter map showing sequencing technologies utilized over the years for given assemblies in Meta-Mined dashboard. *The visualization shows number of assemblies sequenced using specific technologies in given year. The circle size indicates the log count of the assemblies. In the dashboard, this is coupled with dropdown menu that allows users to select assemblies sequenced by one or multiple sequencing technologies*.

**Figure 7.**
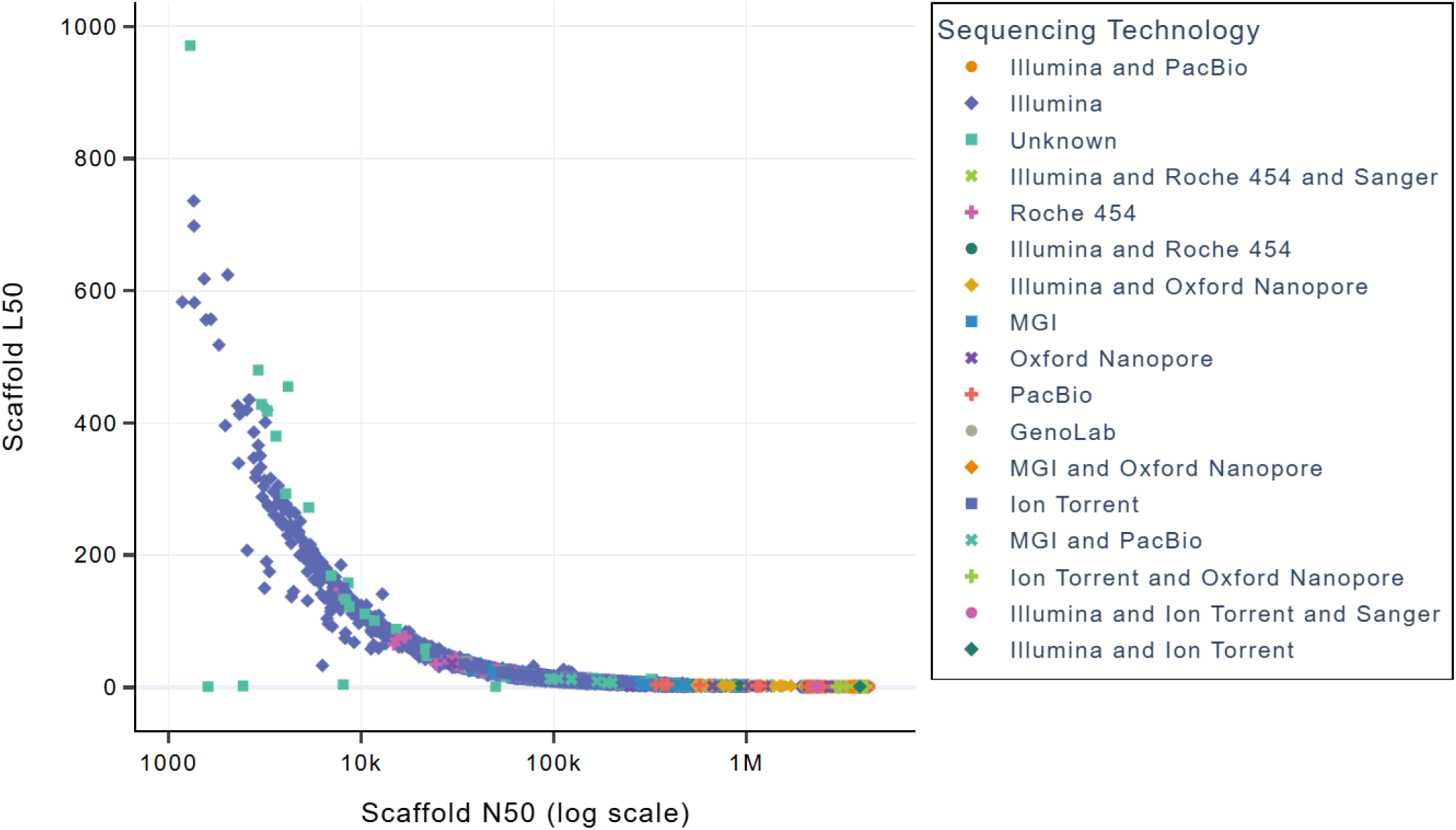
Scatter plot of scaffold N50 vs. L50 values for given assemblies in the Meta-Mined dashboard. *The plot illustrates structural quality of the assemblies in relation with N50 and L50 values. Each point in the plot represents an assembly, with color indicating the sequencing technology used. This visualization is combined with sliders to filter assemblies based-on certain value of L50 and/or N50*.

**Figure 8.**
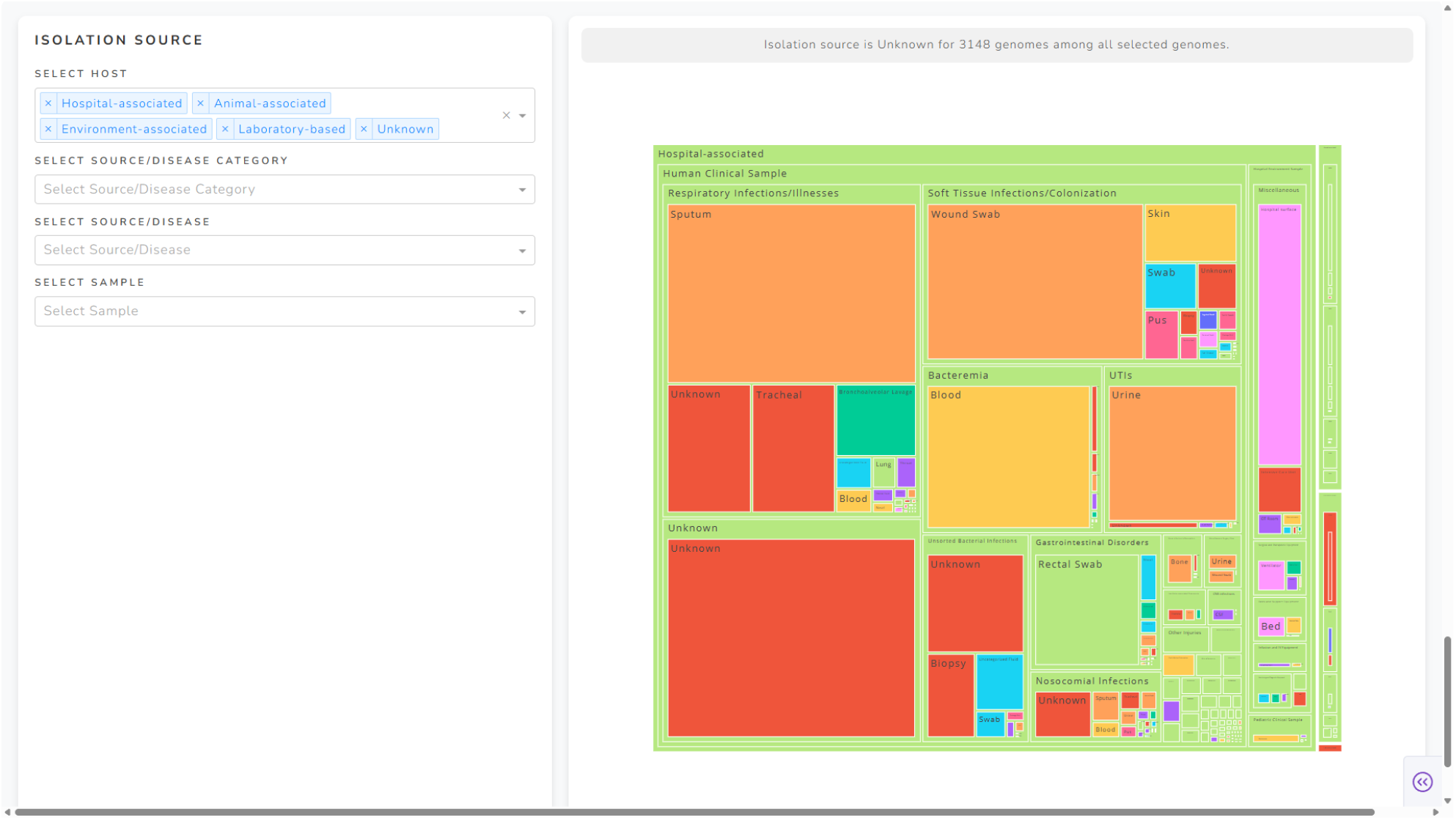
Treemap of normalized isolation sources in the Meta-Mined dashboard. *The image presents a static treemap (on right) showing the category-wise distribution of assemblies within various isolation sources. Each block corresponds to a specific source, with size proportional to its frequency in the dataset. In live dashboard, user can refine the view by selecting or deselecting a certain block and explore the distribution better. The accompanying dropdown filters (on left) further enables users to select the assemblies from one or more isolation sources based-on their need*.

Along with these, the other options allow users to subset assemblies based on annotation provider, assembly level, gene count, BioSample or BioProject IDs, and more. For example, one can retrieve genomes submitted between 2013 and 2024, originating from a specific U.S. state, sequenced using a particular technology, and meeting thresholds such as ANI % identity above 95 and N50 value above 1 Mb, among others. The comprehensive visualizations provided by dashboard makes it possible to tailor data exploration and retrieval to specific research questions. The filtered data then can be used to fetch other genomic data for comparative analysis. The detailed description of the dashboard components is available at https://github.com/prekijpatel/MetaMiner/wiki/Components-of-Dashboard.

## Conclusion

MetaMiner enhances the efficiency of genomic metadata processing by automating data retrieval, normalization, and analysis. Its advanced algorithms standardize diverse metadata fields, enabling comprehensive visualization and exploration. The interactive dashboard allows users to filter and analyse data based on custom criteria, supporting more focused and insightful research. By streamlining the process, MetaMiner improves the accessibility and usability of large-scale genomic datasets, making it an essential tool for data-driven genomic research.

## Declarations

### Availability of data and materials

MetaMiner is available freely at https://github.com/prekijpatel/MetaMiner.

### Author’s contributions

JP conceptualised the study, designed the MetaMiner tool, and performed data curation and analysis. JP also wrote the initial draft of the manuscript. RKE supervised the project, provided critical insights and guided the research framework. RKE also contributed significantly to manuscript revisions and final editing. Both read, reviewed and approved the final manuscript.

## Acknowledgements

JP is supported by the DBT-JRF Fellowship program. The authors sincerely thank all colleagues, particularly Prabhu Balasubramanian sir, Ada Zwetlana, Lily Singh, Ratnabali Ghosh, Sinad Mohamed and Saunak Dasgupta for their valuable time, insightful discussions, and constructive inputs that greatly contributed to the development and refinement of this work.

