## Supplementary material for "MetaMiner: Streamlined GUI Tool for Retrieving, Normalizing and Exploring Metadata": Supplementary File 1.html

Metadata retrieval and transformation by NCBI's datasets and dataformat


### Metadata retrieval and transformation by NCBI's datasets and dataformat

```
Microsoft Windows [Version 10.0.22631.4751]
(c) Microsoft Corporation. All rights reserved.

C:\Users\Prekij>E:

E:\>cd 2025/metaminer/

E:\2025\metaminer>cd "metaminer_bioarchiv"

E:\2025\metaminer\metaminer_bioarchiv>cd abau_comparison

E:\2025\metaminer\metaminer_bioarchiv\abau_comparison>datasets.exe summary genome taxon "Acinetobacter baumannii" --as-json-lines > clid_abau.jsonl

E:\2025\metaminer\metaminer_bioarchiv\abau_comparison>datasets.exe --version
datasets version: 16.40.1

E:\2025\metaminer\metaminer_bioarchiv\abau_comparison>dataformat.exe version
16.40.1

E:\2025\metaminer\metaminer_bioarchiv\abau_comparison>dataformat tsv genome --inputfile ./clid_abau.jsonl > clid_report_transformed_tsv.tsv

E:\2025\metaminer\metaminer_bioarchiv\abau_comparison>dataformat tsv genome --inputfile ./dd_assembly_data_report.jsonl > dd_report_transformed_tsv.tsv

E:\2025\metaminer\metaminer_bioarchiv\abau_comparison>dataformat tsv genome --inputfile ./metaminer_assembly_report.json > metaminer_report_transformed_tsv.tsv

For best results make sure that you're using the latest version of the dataformat command line tool
Download the latest version of the dataformat and datasets command line tools: https://www.ncbi.nlm.nih.gov/datasets/docs/v2/download-and-install

Use --force to remove this warning.

dataformat doesn't recognize this input!
Error: bufio.Scanner: token too long

Use dataformat tsv genome <command> --help for detailed help about a command.


E:\2025\metaminer\metaminer_bioarchiv\abau_comparison>
```

### Metadata retrieval and transformation by MetaMiner

```
2025-02-03 07:56:43,487:INFO - Saving file path selected: E:/2025/metaminer/metaminer bioarchiv/abau_comparison
2025-02-03 07:57:02,715:DEBUG - download_json button pressed!
2025-02-03 07:57:02,715:INFO - Selected to download JSON file.
2025-02-03 07:57:02,715:DEBUG - setting file name to: Acinetobacter_baumannii.json
2025-02-03 07:57:02,715:INFO - Selected organism: prokaryote, Selected genome_gene: genome, Selected accid_taxid: taxon, textbox: "Acinetobacter baumannii"
2025-02-03 07:57:02,715:DEBUG - OS name: nt, datasets executable: datasets.exe
2025-02-03 07:57:02,715:INFO - This command will be executed to download the required JSON file: datasets.exe summary genome taxon "Acinetobacter baumannii" > ./tmp/Acinetobacter_baumannii.json
2025-02-03 07:57:02,715:INFO - Checking for active internet connection.
2025-02-03 07:57:02,778:DEBUG - Active internet connection found.
2025-02-03 07:57:02,778:INFO - Internet connection available. Starting download...
2025-02-03 07:57:02,778:INFO - Downloading metadata. Time elapsed: 0.00 seconds.
2025-02-03 07:57:07,829:INFO - Downloading metadata. Time elapsed: 5.05 seconds.
2025-02-03 07:57:12,843:INFO - Downloading metadata. Time elapsed: 10.06 seconds.
2025-02-03 07:57:17,859:INFO - Downloading metadata. Time elapsed: 15.08 seconds.
2025-02-03 07:57:22,871:INFO - Downloading metadata. Time elapsed: 20.09 seconds.
2025-02-03 07:57:27,875:INFO - Downloading metadata. Time elapsed: 25.10 seconds.
2025-02-03 07:57:32,887:INFO - Downloading metadata. Time elapsed: 30.11 seconds.
2025-02-03 07:57:37,893:INFO - Downloading metadata. Time elapsed: 35.12 seconds.
2025-02-03 07:57:42,907:INFO - Downloading metadata. Time elapsed: 40.13 seconds.
2025-02-03 07:57:47,924:INFO - Downloading metadata. Time elapsed: 45.15 seconds.
2025-02-03 07:57:52,931:INFO - Downloading metadata. Time elapsed: 50.15 seconds.
2025-02-03 07:57:57,938:INFO - Downloading metadata. Time elapsed: 55.16 seconds.
2025-02-03 07:58:02,948:INFO - Downloading metadata. Time elapsed: 60.17 seconds.
2025-02-03 07:58:07,963:INFO - Downloading metadata. Time elapsed: 65.18 seconds.
2025-02-03 07:58:12,977:INFO - Downloading metadata. Time elapsed: 70.20 seconds.
2025-02-03 07:58:17,990:INFO - Downloading metadata. Time elapsed: 75.21 seconds.
2025-02-03 07:58:23,006:INFO - Downloading metadata. Time elapsed: 80.23 seconds.
2025-02-03 07:58:28,013:INFO - Downloading metadata. Time elapsed: 85.24 seconds.
2025-02-03 07:58:30,727:INFO - JSON file downloaded successfully!
2025-02-03 07:58:30,727:INFO - Download complete in 85.24 seconds.
```
