## Supplementary material for "MetaMiner: Streamlined GUI Tool for Retrieving, Normalizing and Exploring Metadata": Supplementary File 2.html

plotting


### Figures and Analysis Code¶

Following script depicts the code used for various analysis and plotting figures for the manuscript.

In [1]:

```
# Importing the necessary libraries/Functions
import pandas as pd
import numpy as np
import plotly.express as px
import logging
import plotly.io as pio
pio.renderers.default = "browser"
from utils import * # the library of functions built for this project and are being used in MetaMiner
import os
import sys
import timeit
```

1. Starting with analyzing the metadata file that has been downloaded using various modes. The data is loaded using the `load_json()` function, which is built to handle `JSON` and `JSONL` file robustly. The loaded `JSON`/`JSONL` is then checked for the records they hold to validate the completeness of the retrieved metadata.

In [2]:

```
# loading data
metaminer_downloaded_json = load_json("./metaminer_bioarchiv/abau_comparison/metaminer_assembly_report.json")
webpage_downloaded_jsonl = load_json("./metaminer_bioarchiv/abau_comparison/dd_assembly_data_report.jsonl")
cmd_downloaded_jsonl = load_json("./metaminer_bioarchiv/abau_comparison/clid_abau.jsonl")
```

In [3]:

```
total_genomes = [webpage_downloaded_jsonl["total_count"], cmd_downloaded_jsonl["total_count"],
                 metaminer_downloaded_json["total_count"]]
print(total_genomes)
```

```
[46806, 46806, 46806]
```

In [4]:

```
# plotting the genome count from each json downloaded using various modalities
figure = px.bar(
    x=["Webpage", "CMD", "MetaMiner"],
    y=total_genomes,
    text=total_genomes,
    color=["Webpage", "CMD", "MetaMiner"],
    color_discrete_sequence= px.colors.qualitative.Pastel
)

figure.update_layout(
    font=dict(size=14, color='black'), 
    bargap=0.3,
    xaxis=dict(
        title="Metadata (json) Source",
        titlefont=dict(size=16, color='black'),
        tickfont=dict(size=14, color='black'),
        # tickangle=-30,
        showline=True,
        linecolor='black',
        ticks='outside',
        tickwidth=2,
        ticklen=6,
    ),
    yaxis=dict(
        title="Metadata for 'n' Assemblies",
        titlefont=dict(size=16, color='black'),
        showline=True,
        linecolor='black',
        tickfont=dict(size=14, color='black'),
        ticks='outside',
        tickwidth=2,
        ticklen=6,
        range = [0, 65000],
        
    ),
    height=500,
    width=440,
    showlegend=False,
    plot_bgcolor="rgba(0, 0, 0, 0)",
    paper_bgcolor="rgba(0,0,0,0)"
)

# figure.write_image("./metaminer_bioarchiv/abau_comparison/total_genomes.png", format="png", scale=4)
```

2. Next, in order to evaluate the formatting efficiency of Metaminer, various parameters were taken into consideration. The formatted files' size, time taken by them to load in current environment, their organization and size.

In [5]:

```
# Calculating the size of formatted data on drive using `os.path` module

sizes_in_bytes = {
    'webpage_downloaded_dataformat_transformed' : os.path.getsize("./metaminer_bioarchiv/abau_comparison/dd_report_transformed_tsv.tsv"),
    'cmd_downloaded_dataformat_transformed' : os.path.getsize("./metaminer_bioarchiv/abau_comparison/clid_report_transformed_tsv.tsv"),
    'metaminer_downloaded_dataformat_transformed' : os.path.getsize("./metaminer_bioarchiv/abau_comparison/metaminer_report_transformed_tsv.tsv"),
    'webpage_downloaded_metaminer_transformed' : os.path.getsize("./metaminer_bioarchiv/abau_comparison/webpage_downloaded_allmetadata_raw.tsv"),
    'cmd_downloaded_metaminer_transformed' : os.path.getsize("./metaminer_bioarchiv/abau_comparison/cmd_downloaded_allmetadata_raw.tsv"),
    'metaminer_downloaded_metaminer_transformed' : os.path.getsize("./metaminer_bioarchiv/abau_comparison/metaminer_downloaded_allmetadata_raw.tsv")
}
```

In [6]:

```
# determining how much time it takes to load the data
def load_clid_report():
    return pd.read_csv("./metaminer_bioarchiv/abau_comparison/clid_report_transformed_tsv.tsv", sep="\t", low_memory=False)
def load_dd_report():
    return pd.read_csv("./metaminer_bioarchiv/abau_comparison/dd_report_transformed_tsv.tsv", sep="\t", low_memory=False)
def load_webpage_downloaded_allmetadata_raw():
    return pd.read_csv("./metaminer_bioarchiv/abau_comparison/webpage_downloaded_allmetadata_raw.tsv", sep="\t", low_memory=False)
def load_cmd_downloaded_allmetadata_raw():
    return pd.read_csv("./metaminer_bioarchiv/abau_comparison/cmd_downloaded_allmetadata_raw.tsv", sep="\t", low_memory=False)
def load_metaminer_downloaded_allmetadata_raw():
    return pd.read_csv("./metaminer_bioarchiv/abau_comparison/metaminer_downloaded_allmetadata_raw.tsv", sep="\t", low_memory=False)


execution_time_clid_report = timeit.timeit(load_clid_report, number=5) # loading them 5 times to get the average time
execution_time_dd_report = timeit.timeit(load_dd_report, number=5)
execution_time_webpage_allmetadata = timeit.timeit(load_webpage_downloaded_allmetadata_raw, number=5)
execution_time_cmd_allmetadata = timeit.timeit(load_cmd_downloaded_allmetadata_raw, number=5)
execution_time_metaminer_allmetadata = timeit.timeit(load_metaminer_downloaded_allmetadata_raw, number=5)


time_taken_to_load_in_seconds = {
    'webpage_downloaded_dataformat_transformed' :  execution_time_dd_report/5,
    'cmd_downloaded_dataformat_transformed' :  execution_time_clid_report/5,
    'webpage_downloaded_metaminer_transformed' : execution_time_webpage_allmetadata/5,
    'cmd_downloaded_metaminer_transformed' : execution_time_cmd_allmetadata/5,
    'metaminer_downloaded_metaminer_transformed' : execution_time_metaminer_allmetadata/5,
}
```

In [7]:

```
# loading the transformed data to know the data structure
cmd_downloaded_dataformat_transformed = pd.read_csv("./metaminer_bioarchiv/abau_comparison/clid_report_transformed_tsv.tsv", sep="\t", low_memory=False)
webpage_downloaded_dataformat_transformed = pd.read_csv("./metaminer_bioarchiv/abau_comparison/dd_report_transformed_tsv.tsv", sep="\t", low_memory=False)
webpage_downloaded_metaminer_transformed = pd.read_csv("./metaminer_bioarchiv/abau_comparison/webpage_downloaded_allmetadata_raw.tsv", sep="\t", low_memory=False)
cmd_downloaded_metaminer_transformed = pd.read_csv("./metaminer_bioarchiv/abau_comparison/cmd_downloaded_allmetadata_raw.tsv", sep="\t", low_memory=False)
metaminer_downloaded_metaminer_transformed = pd.read_csv("./metaminer_bioarchiv/abau_comparison/metaminer_downloaded_allmetadata_raw.tsv", sep="\t", low_memory=False)

shape_of_dataframes = {
    'webpage_downloaded_dataformat_transformed' : webpage_downloaded_dataformat_transformed.shape,
    'cmd_downloaded_dataformat_transformed' : cmd_downloaded_dataformat_transformed.shape,
    'metaminer_downloaded_dataformat_transformed' : (0, 0),
    'webpage_downloaded_metaminer_transformed' : webpage_downloaded_metaminer_transformed.shape,
    'cmd_downloaded_metaminer_transformed' : cmd_downloaded_metaminer_transformed.shape,
    'metaminer_downloaded_metaminer_transformed' : metaminer_downloaded_metaminer_transformed.shape
}
```

In [8]:

```
# putting everything related to transformed data into a dataframe
size_df = pd.DataFrame.from_dict(sizes_in_bytes, orient='index', columns=['Size in Bytes'])
size_df['in_MB'] = (size_df['Size in Bytes'] / (1024 * 1024)).round(3)
size_df['downloaded_from'] = ['Webpage', 'CMD', 'MetaMiner', 'Webpage', 'CMD', 'MetaMiner']
size_df['formatted_using'] = ['Dataformat', 'Dataformat', 'Dataformat', 'MetaMiner', 'MetaMiner', 'MetaMiner']
size_df['time_taken_to_load'] = [42.177, 42.321, 0, 2.876, 2.890, 2.911]
size_df['shape'] = ["(1149789, 174)", "(1149789, 174)", "(0, 0)", "(46806, 353)", "(46806, 353)", "(46806, 353)"]
```

In [9]:

```
size_df
```

Out[9]:

|  | Size in Bytes | in\_MB | downloaded\_from | formatted\_using | time\_taken\_to\_load | shape |
| --- | --- | --- | --- | --- | --- | --- |
| webpage\_downloaded\_dataformat\_transformed | 1657862249 | 1581.061 | Webpage | Dataformat | 42.177 | (1149789, 174) |
| cmd\_downloaded\_dataformat\_transformed | 1657862249 | 1581.061 | CMD | Dataformat | 42.321 | (1149789, 174) |
| metaminer\_downloaded\_dataformat\_transformed | 0 | 0.000 | MetaMiner | Dataformat | 0.000 | (0, 0) |
| webpage\_downloaded\_metaminer\_transformed | 58023174 | 55.335 | Webpage | MetaMiner | 2.876 | (46806, 353) |
| cmd\_downloaded\_metaminer\_transformed | 57929164 | 55.246 | CMD | MetaMiner | 2.890 | (46806, 353) |
| metaminer\_downloaded\_metaminer\_transformed | 57928570 | 55.245 | MetaMiner | MetaMiner | 2.911 | (46806, 353) |

In [10]:

```
# plotting the size and shape of the formatted data with respect to the source of download
#  
fig_s = px.bar(
    size_df,
    x="formatted_using",
    y='in_MB',
    # text='shape',
    color='downloaded_from',
    color_discrete_sequence=px.colors.qualitative.Pastel,
    barmode='group'
)

fig_s.update_layout(
    font=dict(size=14, color='black'), 
    bargap=0.2,
    xaxis=dict(
        title="Formatted using",
        titlefont=dict(size=16, color='black'),
        tickfont=dict(size=14, color='black'),
        # tickangle=-30,
        showline=True,
        linecolor='black',
        ticks='outside',
        tickwidth=2,
        ticklen=6,
    ),
    yaxis=dict(
        title="Size in Megabytes (MB)",
        titlefont=dict(size=16, color='black'),
        showline=True,
        linecolor='black',
        tickfont=dict(size=14, color='black'),
        ticks='outside',
        tickwidth=2,
        ticklen=6,
        range = [0, 1800],
    ),
    height=450,
    width=400,
    showlegend=True,
    plot_bgcolor="rgba(0, 0, 0, 0)",
    paper_bgcolor="rgba(0,0,0,0)"
)

# fig_s.show()
# fig_s.write_image("./metaminer_bioarchiv/abau_comparison/size_shape.png", format="png", scale=4)
```

In [11]:

```
# time taken to load the formatted data
fig = px.bar(
    size_df,
    x="time_taken_to_load",
    y="formatted_using",
    color="downloaded_from",
    color_discrete_sequence=px.colors.qualitative.Pastel,
    text="time_taken_to_load",
    barmode='group',
)

fig.update_layout(
    font=dict(size=14, color='black'), 
    bargap=0.2,
    xaxis=dict(
        title="Time taken to load data (seconds)",
        titlefont=dict(size=16, color='black'),
        tickfont=dict(size=14, color='black'),
        # tickangle=-30,
        showline=True,
        linecolor='black',
        ticks='outside',
        tickwidth=2,
        ticklen=6,
        range=[0, 50]
    ),
    yaxis=dict(
        title="Formatted using",
        titlefont=dict(size=16, color='black'),
        showline=True,
        linecolor='black',
        tickfont=dict(size=14, color='black'),
        ticks='outside',
        tickwidth=2,
        ticklen=6,
        # range = [0, 65000],
        
    ),
    height=350,
    width=500,
    showlegend=False,
    plot_bgcolor="rgba(0, 0, 0, 0)",
    paper_bgcolor="rgba(0,0,0,0)"
)

# fig.write_image("./metaminer_bioarchiv/abau_comparison/time_to_load.png", format="png", scale=4)
```

3. Next, to validate the normalization efficiency, We again used the MetaMiner downloaded and transformed metadata of *Acinetobacter baumannii*. However, to make validation feasible, we randomly selected 2500 genomes out of the raw transformed data. Then, we compared the MetaMiner-based normalization with Manual Curation.

In [12]:

```
# Loading the Raw metadata
Raw_trasnformed = pd.read_csv("./metaminer_bioarchiv/abau_comparison/metaminer_downloaded_allmetadata_raw.tsv", sep="\t", low_memory=False)
```

In [13]:

```
# randomly selecting 2500 assemblies' metadata  from the raw metadata

randomly_selected_assemblies = Raw_trasnformed.sample(n=2500, random_state=801)
randomly_selected_assemblies.shape
```

Out[13]:

```
(2500, 353)
```

In [14]:

```
# preparing and saving the data for manual curation
unique_locations = pd.Series(randomly_selected_assemblies['geo_loc_name'].unique(), name='geo_loc_name')
unique_isolation_samples = randomly_selected_assemblies[['host', 'host_disease', 'isolation_source']].drop_duplicates()
unique_sequencing_technology = pd.Series(randomly_selected_assemblies['Sequencing_Technology'].unique(), name='Sequencing_Technology')

len(unique_locations), len(unique_isolation_samples), len(unique_sequencing_technology)
# unique_locations.to_csv("./metaminer_bioarchiv/abau_comparison/unique_locations.tsv", sep="\t", index=False)
# unique_isolation_sources.to_csv("./metaminer_bioarchiv/abau_comparison/unique_isolation_sources.tsv", sep="\t", index=False)
# unique_sequencing_technology.to_csv("./metaminer_bioarchiv/abau_comparison/unique_sequencing_technology.tsv", sep="\t", index=False)
```

Out[14]:

```
(339, 740, 79)
```

Post saving this, the saved `TSVs` were manually curated and the curated data was loaded back to the notebook for further analysis. Starting With:

- Geographical Location Normalization:

In [15]:

```
# normalizing the data using MetaMiner
normalized_cleaned_data = all_normalization_operations(randomly_selected_assemblies)
normalized_cleaned_data.to_csv("./metaminer_bioarchiv/abau_comparison/normalized_cleaned_data.tsv", sep="\t", index=False)
normalized_cleaned_data.shape
```

Out[15]:

```
(2500, 366)
```

In [16]:

```
# loading the manually curated data and mapping the same into dataframe

## loading all

manually_curated_locations = pd.read_csv("./metaminer_bioarchiv/abau_comparison/unique_locations_sorted.tsv", sep="\t", encoding='iso-8859-1')
manually_curated_isolation_sources = pd.read_csv("./metaminer_bioarchiv/abau_comparison/unique_isolation_sources_sorted_1.tsv", sep='\t', low_memory=False,)
manually_curated_isolation_sources = manually_curated_isolation_sources.where(pd.notnull(manually_curated_isolation_sources),None)
manually_curated_sequencing_technology_df = pd.read_csv("./metaminer_bioarchiv/abau_comparison/unique_sequencing_technology_sorted.tsv", sep='\t', low_memory=False)


## Mapping geographical data into dataframe
manual_country_state_map = {row[0]: (row[1], row[2], row[3], row[4], row[5]) for row in manually_curated_locations.itertuples(index=False)}

### mapping the curated data to dataframe where MetaMiner curated data is also there
normalized_cleaned_data['Manual_country_name'] = normalized_cleaned_data['geo_loc_name'].map(
    lambda x: manual_country_state_map.get(x, (None, None, None, None, None))[0] if x != None else None)
normalized_cleaned_data['Manual_country_common_name'] = normalized_cleaned_data['geo_loc_name'].map(
    lambda x: manual_country_state_map.get(x, (None, None, None, None, None))[1] if x != None else None)
normalized_cleaned_data['Manual_country_three_lettered_name'] = normalized_cleaned_data['geo_loc_name'].map(
    lambda x: manual_country_state_map.get(x, (None, None, None, None, None))[2] if x != None else None)
normalized_cleaned_data['Manual_state_name'] = normalized_cleaned_data['geo_loc_name'].map(
    lambda x: manual_country_state_map.get(x, (None, None, None, None, None))[3] if x != None else None)
normalized_cleaned_data['Manual_state_code'] = normalized_cleaned_data['geo_loc_name'].map(
    lambda x: manual_country_state_map.get(x, (None, None, None, None, None))[4] if x != None else None)

print(normalized_cleaned_data.shape)

## Mapping curated Isolation sources 
normalized_cleaned_data[['host', 'host_disease', 'isolation_source', 'identified_host', 'source_category', 'source_y', 'sample']] = normalized_cleaned_data[['host', 'host_disease', 'isolation_source', 'identified_host', 'source_category', 'source_y', 'sample']].where(pd.notnull(normalized_cleaned_data[['host', 'host_disease', 'isolation_source', 'identified_host', 'source_category', 'source_y', 'sample']]),None)

normalized_cleaned_data = normalized_cleaned_data.merge(manually_curated_isolation_sources, how='left', on=['host', 'host_disease', 'isolation_source'])

normalized_cleaned_data.drop_duplicates(inplace=True)
normalized_cleaned_data.reset_index(drop=True, inplace=True)
print(normalized_cleaned_data.shape)

## Mapping curated Sequencing Technology data 
manually_curated_sequencing_technology_dict = {}

for i in range(len(manually_curated_sequencing_technology_df)):
    key = manually_curated_sequencing_technology_df.iloc[i, 0]
    value = manually_curated_sequencing_technology_df.iloc[i, 1]  
    manually_curated_sequencing_technology_dict[key] = value
                                                
### mapping the manual entries to raw values
normalized_cleaned_data['manually_curated_sequencing_technology'] = normalized_cleaned_data['Sequencing_Technology'].map(manually_curated_sequencing_technology_dict)

print(normalized_cleaned_data.shape)
```

```
(2500, 371)
(2500, 375)
(2500, 376)
```

In [17]:

```
# Comparing Country IDs from MetaMiner and manually curated data as they are the ones which are used for filtering and plotting ultimately

## to avoid differences between nan and none

normalized_cleaned_data[['geo_loc_name', 'country_three_lettered_name', 'Manual_country_three_lettered_name']] = normalized_cleaned_data[['geo_loc_name', 'country_three_lettered_name', 'Manual_country_three_lettered_name']].where(pd.notnull(normalized_cleaned_data[['geo_loc_name', 'country_three_lettered_name', 'Manual_country_three_lettered_name']]),None)


normalization_didt_match = []
metaminer_standardized = []
manually_standardized = []


for i in range(len(normalized_cleaned_data)):
    if normalized_cleaned_data.loc[i, 'country_three_lettered_name'] == normalized_cleaned_data.loc[i, 'Manual_country_three_lettered_name']:
        pass
    else:
        normalization_didt_match.append(normalized_cleaned_data.loc[i, 'geo_loc_name'])
        metaminer_standardized.append([normalized_cleaned_data.loc[i, 'country_name'], normalized_cleaned_data.loc[i, 'country_three_lettered_name']])
        manually_standardized.append([normalized_cleaned_data.loc[i, 'Manual_country_name'] ,normalized_cleaned_data.loc[i, 'Manual_country_three_lettered_name']])

print(f"Not matched entries: {normalization_didt_match}")
print(f"Metaminer curated entries for non-matched values: {metaminer_standardized}")
print(f"Manual curated entries for non-matched values: {manually_standardized}")

total_entries = len(pd.notnull(normalized_cleaned_data['geo_loc_name']))
Total_didnt_match = len(normalization_didt_match)

print(f"Total entries: {total_entries}")
print(f"Total entries that didn't match: {Total_didnt_match}")
```

```
Not matched entries: ['Kosovo:Pristina', 'Korea: Seongnam', 'France:Tahiti', 'Korea: Seongnam']
Metaminer curated entries for non-matched values: [['Serbia', 'SRB'], ["Korea, Democratic People's Republic of", 'PRK'], ['France', 'FRA'], ["Korea, Democratic People's Republic of", 'PRK']]
Manual curated entries for non-matched values: [['Kosovo', 'XKX'], ['Korea, Republic of', 'KOR'], ['French Polynesia', 'PYF'], ['Korea, Republic of', 'KOR']]
Total entries: 2500
Total entries that didn't match: 4
```

In [18]:

```
# Comparing States IDs next

## to avoid differences between nan and none

normalized_cleaned_data[['geo_loc_name', 'state_code', 'Manual_state_code']] = normalized_cleaned_data[['geo_loc_name', 'state_code', 'Manual_state_code']].where(pd.notnull(normalized_cleaned_data[['geo_loc_name', 'state_code', 'Manual_state_code']]),None)


normalization_didt_match = []
metaminer_standardized = []
manually_standardized = []


for i in range(len(normalized_cleaned_data)):
    if normalized_cleaned_data.loc[i, 'state_code'] == normalized_cleaned_data.loc[i, 'Manual_state_code']:
        pass
    else:
        normalization_didt_match.append(normalized_cleaned_data.loc[i, 'geo_loc_name'])
        metaminer_standardized.append([normalized_cleaned_data.loc[i, 'state_name'], normalized_cleaned_data.loc[i, 'state_code']])
        manually_standardized.append([normalized_cleaned_data.loc[i, 'Manual_state_name'] ,normalized_cleaned_data.loc[i, 'Manual_state_code']])

print(f"Not matched entries: {normalization_didt_match}")
print(f"Metaminer curated entries for non-matched values: {metaminer_standardized}")
print(f"Manual curated entries for non-matched values: {manually_standardized}")

total_entries = len(pd.notnull(normalized_cleaned_data['geo_loc_name']))
Total_didnt_match = len(normalization_didt_match)

print(f"Total entries: {total_entries}")
print(f"Total entries that didn't match: {Total_didnt_match}")
```

```
Not matched entries: ['Mexico:Guadalajara', 'Mexico:Guadalajara', 'Mexico:Guadalajara', 'Korea: Seongnam', 'Mexico:Guadalajara', 'France:Tahiti', 'Mexico:Guadalajara', 'Korea: Seongnam', 'Mexico:Guadalajara', 'Mexico:Guadalajara', 'Switzerland: Basel']
Metaminer curated entries for non-matched values: [['MÃ©xico', 'MX-MEX'], ['MÃ©xico', 'MX-MEX'], ['MÃ©xico', 'MX-MEX'], [nan, None], ['MÃ©xico', 'MX-MEX'], [nan, None], ['MÃ©xico', 'MX-MEX'], [nan, None], ['MÃ©xico', 'MX-MEX'], ['MÃ©xico', 'MX-MEX'], ['Basel-Landschaft', 'CH-BL']]
Manual curated entries for non-matched values: [['Jalisco', 'MX-JAL'], ['Jalisco', 'MX-JAL'], ['Jalisco', 'MX-JAL'], ['Gyeonggi Province', 'KR-41'], ['Jalisco', 'MX-JAL'], ['Winward Islands', 'PF-WI'], ['Jalisco', 'MX-JAL'], ['Gyeonggi Province', 'KR-41'], ['Jalisco', 'MX-JAL'], ['Jalisco', 'MX-JAL'], ['Basel-Stadt', 'CH-BS']]
Total entries: 2500
Total entries that didn't match: 11
```

In [19]:

```
# plotting 

match_df = pd.DataFrame({
    'Category': ['Country', 'State'],
    'Matched': [2496, 2489],
})

matrix = match_df.set_index('Category').T 

fig = px.imshow(matrix, 
        labels=dict(x="Category", y="Method", color="Count"),
        x=matrix.columns, y=matrix.index, 
        color_continuous_scale='YlGnBu',
        text_auto=True)

fig.update_layout(
    font=dict(size=14, color='black'), 
    xaxis=dict(
        title="Geographical Location",
        titlefont=dict(size=16, color='black'),
        tickfont=dict(size=14, color='black'),
        showline=True,
        linecolor='black',
        ticks='outside',
        tickangle = -45,
        tickwidth=2,
        ticklen=6,
    ),
    yaxis=dict(
        title = None,
        tickfont=dict(size=14, color='black'),
        showline=True,
        linecolor='black',
        ticks='outside',
        tickwidth=2,
        ticklen=6,
    ),
    height=450,
    width=500,
    showlegend=False,  
    plot_bgcolor="rgba(0, 0, 0, 0)",
    paper_bgcolor="rgba(0, 0, 0, 0)",
    coloraxis_showscale=False
)

# fig.show()

fig.write_image("./metaminer_bioarchiv/abau_comparison/geo_loc_normalized.png", format="png", scale=4)
```

- Isolation sources for organism:

In [20]:

```
# Comparing the isolation source normalization - with and without database supported and database non-supported metaminer curation

## running the all normalization without suppoting database

fuzzy_to_isolation_source_map = {} # defining new dict for normalization, usually this is where the database is loaded

just_the_isolation_sources = randomly_selected_assemblies[['host', 'host_disease', 'isolation_source']].drop_duplicates()

updated_df_isolation_sources_categorized = dict_update_from_new_isolation_sources(just_the_isolation_sources, fuzzy_to_isolation_source_map)

print(updated_df_isolation_sources_categorized.columns)

updated_df_isolation_sources_categorized.rename(columns={'identified_host': 'identified_host_wo_d', 'source_category': 'source_category_wo_d', 'source':'source_wo_d', 'sample':'sample_d'}, inplace=True)

print(updated_df_isolation_sources_categorized.columns)

print(normalized_cleaned_data.shape)


## mapping the things back to the dataframe
normalized_cleaned_data = map_isolation_sources(normalized_cleaned_data, map_df=updated_df_isolation_sources_categorized)
print(normalized_cleaned_data.shape)
```

```
Index(['host', 'host_disease', 'isolation_source', 'identified_host',
       'source_category', 'source', 'sample'],
      dtype='object')
Index(['host', 'host_disease', 'isolation_source', 'identified_host_wo_d',
       'source_category_wo_d', 'source_wo_d', 'sample_d'],
      dtype='object')
(2500, 376)
(2500, 380)
```

In [21]:

```
# first comparing for identified host

host_submitted_entry = []
host_submitted_entry_wo_d = []
host_metaminer = []
host_metaminer_wo_d = []
host_manual = []
host_manual_wo_d = []

number = {}
for i in range(len(normalized_cleaned_data)):
    if normalized_cleaned_data.loc[i,'identified_host'] == normalized_cleaned_data.loc[i,'Manual_identified_host']:
        pass
    else:
        host_submitted_entry.append(normalized_cleaned_data.loc[i,'host'])
        host_metaminer.append(normalized_cleaned_data.loc[i,'identified_host'])
        host_manual.append(normalized_cleaned_data.loc[i,'Manual_identified_host'])

        key = (normalized_cleaned_data.loc[i, 'host'],
               normalized_cleaned_data.loc[i, 'host_disease'],
               normalized_cleaned_data.loc[i, 'isolation_source'])

        value = [normalized_cleaned_data.loc[i, 'identified_host'], 
                 normalized_cleaned_data.loc[i, 'Manual_identified_host']]
        
        number[key] = value

number1 = {}
for i in range(len(normalized_cleaned_data)):
    if normalized_cleaned_data.loc[i,'identified_host_wo_d'] == normalized_cleaned_data.loc[i,'Manual_identified_host']:
        pass
    else:
        host_submitted_entry_wo_d.append(normalized_cleaned_data.loc[i,'host'])
        host_metaminer_wo_d.append(normalized_cleaned_data.loc[i,'identified_host_wo_d'])
        host_manual_wo_d.append(normalized_cleaned_data.loc[i,'Manual_identified_host'])

        key = (normalized_cleaned_data.loc[i, 'host'],
               normalized_cleaned_data.loc[i, 'host_disease'],
               normalized_cleaned_data.loc[i, 'isolation_source'])

        value = [normalized_cleaned_data.loc[i, 'identified_host_wo_d'], 
                 normalized_cleaned_data.loc[i, 'Manual_identified_host']]
        
        number1[key] = value
        

print(f"Host submitted entry: {host_submitted_entry}")
print(f"Host metaminer curated with database: {host_metaminer}")
print(f"Host manually curated: {host_manual}")

print(f"Host submitted entry without databse: {host_submitted_entry_wo_d}")
print(f"Host metaminer curated without databse: {host_metaminer_wo_d}")
print(f"Host manually curated: {host_manual_wo_d}")

total_entries = len(pd.notnull(normalized_cleaned_data['host']))
Total_didnt_match = len(host_submitted_entry)
Total_didnt_match_wo_d = len(host_metaminer_wo_d)

print(f"Total entries: {total_entries}")
print(f"Total entries that didn't match when the metaminer curation was supported by database: {Total_didnt_match}")
print(f"Total entries that didn't match when the metaminer curation was not supported by database: {Total_didnt_match_wo_d}")
print(number)
print(number1)
```

```
Host submitted entry: []
Host metaminer curated with database: []
Host manually curated: []
Host submitted entry without databse: [None, None, None, None, None, None]
Host metaminer curated without databse: ['Animal-associated', 'Hospital-associated', 'Hospital-associated', 'Environment-associated', 'Hospital-associated', 'Unknown']
Host manually curated: ['Unknown', 'Unknown', 'Unknown', 'Unknown', 'Unknown', 'Hospital-associated']
Total entries: 2500
Total entries that didn't match when the metaminer curation was supported by database: 0
Total entries that didn't match when the metaminer curation was not supported by database: 6
{}
{(None, None, 'oral swab'): ['Animal-associated', 'Unknown'], (None, None, 'respiratory'): ['Hospital-associated', 'Unknown'], (None, None, 'soil/water/rectum'): ['Environment-associated', 'Unknown'], (None, None, "the outer surface of the doctor's overall"): ['Unknown', 'Hospital-associated']}
```

In [22]:

```
# now trying with host disease/source category


source_category_submitted_entry = []
source_category_submitted_entry_wo_d = []
source_category_metaminer = []
source_category_metaminer_wo_d = []
source_category_manual = []
source_category_manual_wo_d = []


for i in range(len(normalized_cleaned_data)):
    if normalized_cleaned_data.loc[i,'source_category'] == normalized_cleaned_data.loc[i,'Manual_source_category']:
        pass
    else:
        source_category_submitted_entry.append(normalized_cleaned_data.loc[i,'host_disease'])
        source_category_metaminer.append(normalized_cleaned_data.loc[i,'source_category'])
        source_category_manual.append(normalized_cleaned_data.loc[i,'Manual_source_category'])

number1 = {}
for i in range(len(normalized_cleaned_data)):
    if normalized_cleaned_data.loc[i,'source_category_wo_d'] == normalized_cleaned_data.loc[i,'Manual_source_category']:
        pass
    else:
        source_category_submitted_entry_wo_d.append(normalized_cleaned_data.loc[i,'host_disease'])
        source_category_metaminer_wo_d.append(normalized_cleaned_data.loc[i,'source_category_wo_d'])
        source_category_manual_wo_d.append(normalized_cleaned_data.loc[i,'Manual_source_category'])
        key = (normalized_cleaned_data.loc[i, 'host'],
               normalized_cleaned_data.loc[i, 'host_disease'],
               normalized_cleaned_data.loc[i, 'isolation_source'])

        value = [normalized_cleaned_data.loc[i, 'source_category_wo_d'], 
                 normalized_cleaned_data.loc[i, 'Manual_source_category']]
        
        number1[key] = value
        

print(f"Source Category submitted entry: {source_category_submitted_entry}")
print(f"Source category - metaminer curated with database: {source_category_metaminer}")
print(f"Source category - manually curated: {source_category_manual}")

print(f"Source category submitted entry without database: {source_category_submitted_entry_wo_d}")
print(f"Source category - metaminer curated without database: {source_category_metaminer_wo_d}")
print(f"Source category - manually curated: {source_category_manual_wo_d}")

total_entries = len(pd.notnull(normalized_cleaned_data['host_disease']))
Total_didnt_match = len(source_category_metaminer)
Total_didnt_match_wo_d = len(source_category_metaminer_wo_d)

print(f"Total entries: {total_entries}")
print(f"Total entries that didn't match when the metaminer curation was supported by database: {Total_didnt_match}")
print(f"Total entries that didn't match when the metaminer curation was not supported by database: {Total_didnt_match_wo_d}")

print(number1)
```

```
Source Category submitted entry: ['a. baumannii bacteremia (catheter-realated infection)', 'infection']
Source category - metaminer curated with database: ['Hospital Environment Sample', 'Human Clinical Sample']
Source category - manually curated: ['Human Clinical Sample', 'Hospital Environment Sample']
Source category submitted entry without database: ['a. baumannii bacteremia (catheter-realated infection)', 'urinary tract infection', None, 'pneumonia', None, None, None, None, 'bloodstream infection', 'unknown', None, None, 'nicu', 'pneumonia', 'infection', None, None, None, None, 'infection', None, 'surveillance', 'acinetobacter infection', 'endotracheal infection', None]
Source category - metaminer curated without database: ['Hospital Environment Sample', 'Human Clinical Sample', 'Human Clinical Sample', 'Human Clinical Sample', 'Unknown', 'Hospital Environment Sample', 'Unknown', 'Human Clinical Sample', 'Human Clinical Sample', 'Human Clinical Sample', 'Human Clinical Sample', 'Unknown', 'Hospital Environment Sample', 'Human Clinical Sample', 'Human Clinical Sample', 'Unknown', 'Unknown', 'Unknown', 'Unknown', 'Human Clinical Sample', 'Plant', 'Hospital Environment Sample', 'Human Clinical Sample', 'Human Clinical Sample', 'Human Clinical Sample']
Source category - manually curated: ['Human Clinical Sample', 'Hospital Environment Sample', 'Unknown', 'Hospital Environment Sample', 'Food', 'Human Clinical Sample', 'Food', 'Unknown', 'Hospital Environment Sample', 'Hospital Environment Sample', 'Unknown', 'Food', 'Human Clinical Sample', 'Hospital Environment Sample', 'Hospital Environment Sample', 'Hospital Environment Sample', 'Food', 'Food', 'Food', 'Hospital Environment Sample', 'Food', 'Human Clinical Sample', 'Hospital Environment Sample', 'Hospital Environment Sample', 'Hospital Environment Sample']
Total entries: 2500
Total entries that didn't match when the metaminer curation was supported by database: 2
Total entries that didn't match when the metaminer curation was not supported by database: 25
{('homo sapiens', 'a. baumannii bacteremia (catheter-realated infection)', 'seoul national university bundang hospital intensive care unit'): ['Hospital Environment Sample', 'Human Clinical Sample'], ('homo sapiens', 'urinary tract infection', 'urinary catheter'): ['Human Clinical Sample', 'Hospital Environment Sample'], (None, None, 'respiratory'): ['Human Clinical Sample', 'Unknown'], ('homo sapiens', 'pneumonia', 'hospital environment'): ['Human Clinical Sample', 'Hospital Environment Sample'], (None, None, 'food'): ['Unknown', 'Food'], ('homo sapiens', None, 'catheter urine'): ['Hospital Environment Sample', 'Human Clinical Sample'], ('homo sapiens', 'bloodstream infection', 'foley catheter'): ['Human Clinical Sample', 'Hospital Environment Sample'], ('homo sapiens', 'unknown', 'central venous access devices'): ['Human Clinical Sample', 'Hospital Environment Sample'], ('homo sapiens', 'nicu', 'hand'): ['Hospital Environment Sample', 'Human Clinical Sample'], ('homo sapiens', 'pneumonia', 'drainage basin water'): ['Human Clinical Sample', 'Hospital Environment Sample'], ('homo sapiens', 'infection', 'trachy'): ['Human Clinical Sample', 'Hospital Environment Sample'], (None, None, "the outer surface of the doctor's overall"): ['Unknown', 'Hospital Environment Sample'], ('homo sapiens', 'infection', 'tip of intravascular catheter'): ['Human Clinical Sample', 'Hospital Environment Sample'], (None, None, 'coconut meat [foodon:00003856]; ready-to-eat (rte) [foodon:03316636]; food (chunks) [foodon:00004555]; food (frozen) [foodon:03302148]; retail environment [envo:01001448]'): ['Plant', 'Food'], ('homo sapiens', 'surveillance', 'rectal swab'): ['Hospital Environment Sample', 'Human Clinical Sample'], ('homo sapiens', 'acinetobacter infection', 'endobronchial tube'): ['Human Clinical Sample', 'Hospital Environment Sample'], ('homo sapiens', 'endotracheal infection', 'endotracheal tube'): ['Human Clinical Sample', 'Hospital Environment Sample'], (None, None, 'crash trolley 2_na_room 7'): ['Human Clinical Sample', 'Hospital Environment Sample']}
```

In [23]:

```
# next, the actual source

source_entry = []
source_entry_wo_d = []
source_metaminer = []
source_metaminer_wo_d = []
source_manual = []
source_manual_wo_d = []

number = {}
for i in range(len(normalized_cleaned_data)):
    if normalized_cleaned_data.loc[i,'source_y'] == normalized_cleaned_data.loc[i,'Manual_source_y']:
        pass
    else:
        source_entry.append(normalized_cleaned_data.loc[i,'isolation_source'])
        source_metaminer.append(normalized_cleaned_data.loc[i,'source_y'])
        source_manual.append(normalized_cleaned_data.loc[i,'Manual_source_y'])
        key = (normalized_cleaned_data.loc[i, 'host'],
               normalized_cleaned_data.loc[i, 'host_disease'],
               normalized_cleaned_data.loc[i, 'isolation_source'])

        value = [normalized_cleaned_data.loc[i, 'source_y'], 
                 normalized_cleaned_data.loc[i, 'Manual_source_y']]
        
        number[key] = value

number1 = {}
for i in range(len(normalized_cleaned_data)):
    if normalized_cleaned_data.loc[i,'source_wo_d'] == normalized_cleaned_data.loc[i,'Manual_source_y']:
        pass
    else:
        source_entry_wo_d.append(normalized_cleaned_data.loc[i,'isolation_source'])
        source_metaminer_wo_d.append(normalized_cleaned_data.loc[i,'source_wo_d'])
        source_manual_wo_d.append(normalized_cleaned_data.loc[i,'Manual_source_y'])
        key = (normalized_cleaned_data.loc[i, 'host'],
               normalized_cleaned_data.loc[i, 'host_disease'],
               normalized_cleaned_data.loc[i, 'isolation_source'])

        value = [normalized_cleaned_data.loc[i, 'source_wo_d'], 
                 normalized_cleaned_data.loc[i, 'Manual_source_y']]
        
        number1[key] = value


print(f"Source submitted entry: {source_entry}")
print(f"Source - metaminer curated with database: {source_metaminer}")
print(f"Source - manually curated: {source_manual}")

print(f"Source submitted entry without database: {source_entry_wo_d}")
print(f"Source - metaminer curated without database: {source_metaminer_wo_d}")
print(f"Source - manually curated: {source_manual_wo_d}")

total_entries = len(pd.notnull(normalized_cleaned_data['isolation_source']))
Total_didnt_match = len(source_metaminer)
Total_didnt_match_wo_d = len(source_metaminer_wo_d)

print(f"Total entries: {total_entries}")
print(f"Total entries that didn't match when the metaminer curation was supported by database: {Total_didnt_match}")
print(f"Total entries that didn't match when the metaminer curation was not supported by database: {Total_didnt_match_wo_d}")

print(number)
print(number1)
```

```
Source submitted entry: ['seoul national university bundang hospital intensive care unit', 'trachy']
Source - metaminer curated with database: ['Infusion and IV Equipment', 'Unsorted Bacterial infections']
Source - manually curated: ['Bacteremia', 'Surgical and Therapeutic Equipment']
Source submitted entry without database: ['lavage fluid', 'trachea', 'anal margin', 'blood', 'lavage fluid', 'swab of axilla and groin', 'trach asp', 'rectal', 'tracheal aspirate', 'sputum', 'rectal', 'bronchial wash', 'tracheal suction', 'seoul national university bundang hospital intensive care unit', 'tracheal aspirate', 'bronchial wash', 'periprosthetic liquid', 'c tra asp', 'abdomen isolate', 'swab_wound', 'urinary catheter', 'wash_bronchoalveolar lavage (bal)', 'tracheal secretion', 'respiratory', 'respiratory', 'tracheal swab', 'axilla/groin swab', 'endotracheal aspirate', 'tracheal aspirate isolate', 'tracheal secretion', 'swab (wound)', 'tracheal aspirate', 'hospital environment', 'tracheal aspirate', 'tracheal aspirate', 'sputum', 'sputum', 'lung', 'catheter urine', 'axilla/groin swab', 'axilla and groin swab', 'wound isolate', 'endotracheal aspirate', 'background grass (no manure on field)', 'rectal', 'broncho-alveolar lavage', 'rectal', 'wound isolate', 'skin; arms/legs', 'axilla/groin', 'respiratory', 'foley catheter', 'tracheal secretion', 'respiratory', 'central venous access devices', 'respiratory', 'background grass (no manure on field)', 'lung', 'endotracheal aspirate', 'bronchial wash', 'uroculture', 'sputum', 'bone', 'urine', 'trach asp', 'urine', 'axilla and groin swab', 'burn patient', 'axilla and groin swab', 'tracheal aspirate', 'wash_bronchoalveolar lavage (bal)', 'bedside rail in hospital intensive care unit', 'tracheal secretion', 'abdominal wound isolate', 'sputum', 'hydrothorax', 'sputum', 'secreta', 'etta', 'penis swab', 'rectal', 'tracheal aspirate', 'sput', 'trach asp', 'wound isolate', 'tracheal secretion', 'right shin blister', 'hand', 'tracheal aspirate', 'swab', 'drainage basin water', 'trachy', 'blood isolate', 'oral swab_p101_bu22', 'etta', 'urine', 'tracheal secretion', 'urethral swab', 'respiratory', 'low respiratory tract', 'sputum', 'axilla and groin swab', 'rectal', 'sputum', 'urine', 'tracheal aspirate', 'rectal', 'foot', 'sputum', 'oral swab_p23_bu25', 'rectal', 'heel', 'respiratory', 'respiratory', 'trachael aspirate', 'rectal', 'tip of intravascular catheter', 'rectal', 'blood', 'tracheal aspirate', 'coconut meat [foodon:00003856]; ready-to-eat (rte) [foodon:03316636]; food (chunks) [foodon:00004555]; food (frozen) [foodon:03302148]; retail environment [envo:01001448]', 'missing', 'hip', 'trach asp', 'blood isolate', 'tracheal secretion', 'rectal', 'axilla/groin swab', 'swab_wound', 'sputum', 'rectal', 'tracheal secretion', 'sputum', 'rectal swab', 'tissue;sacrum', 'ett', 'trachea', 'respiratory', 'tracheal aspirate/wash', 'respiratory', 'urine', 'tracheal aspirate', 'sputum', 'swab (wound)', 'tracheal aspirate isolate', 'wound isolate', 'severe trauma site', 'urine', 'rectal', 'sputum', 'tracheal aspirate', 'tracheal secretion', 'draining liquid of kidney transplantation recipient', 'sputum', 'rectal swab_p78_bu13', 'bronch wash', 'trach asp', 'perirectal', 'axilla/groin swab', 'endobronchial tube', 'sputum', 'tracheal aspirate', 'endotracheal tube', 'endotracheal aspirate', 'axilla and groin swab', 'left hip', 'tracheal aspirate', 'tracheal aspirate', 'endotracheal aspirate', 'abdomen', 'sputum', 'tracheal aspirate', 'tracheal secretions', 'trach asp', 'other isolate', 'respiratory', 'urine', 'tracheal aspirate', 'endotracheal aspirate', 'leg, left', 'respiratory', 'tracheal aspirate', 'tissue_lung', 'blood isolate', 'control grass (no manure)', 'trachea', 'skin damage', 'skin; arms and legs', 'rectal', 'lung', 'pleural effusion', 'tracheal aspirate/wash', 'tracheal aspirate/wash', 'crash trolley 2_na_room 7', 'rectal', 'scrotal', 'low respiratory tract', 'st', 'rectal swab_p26_bu28', 'oral swab_p96_bu15', 'axilla/groin swab', 'axilla and groin swab', None, 'tracheal aspirate', 'chest', 'blood', 'urine', 'sputamentum', 'tracheal secretion', 'tracheal aspirate', 'throat wash', 'perirectal swab']
Source - metaminer curated without database: ['Gastrointestinal Disorders', 'Unknown', 'Unknown', 'Bacteremia', 'Gastrointestinal Disorders', 'Unknown', 'Unknown', 'Unknown', 'Unknown', 'RTIs', 'Unknown', 'Allergic Disorders', 'Unknown', 'Infusion and IV Equipment', 'Unknown', 'Allergic Disorders', 'Unknown', 'Unknown', 'Bone Infection', 'Other Injuries', 'UTIs', 'Immunodeficiency Disorders', 'Unknown', 'RTIs', 'Unsorted Bacterial infections', 'Unknown', 'Unknown', 'Unknown', 'Unknown', 'Unknown', 'Other Injuries', 'Unknown', 'RTIs', 'Unknown', 'Unknown', 'RTIs', 'RTIs', 'Unknown', 'Infusion and IV Equipment', 'Unknown', 'Unknown', 'Bone Infection', 'Unknown', 'Agricultural Soil', 'Unsorted Bacterial infections', 'RTIs', 'Unknown', 'Bone Infection', 'Unknown', 'Unknown', 'RTIs', 'Bacteremia', 'Unknown', 'Unsorted Bacterial infections', 'Unknown', 'RTIs', 'Agricultural Soil', 'Unknown', 'Unknown', 'Allergic Disorders', 'Soft Tissue Infections/Colonization', 'RTIs', 'Bone Infection', 'Soft Tissue Infections/Colonization', 'Unknown', 'Soft Tissue Infections/Colonization', 'Unknown', 'CNS infections', 'Unknown', 'Unknown', 'Immunodeficiency Disorders', 'Miscellaneous', 'Unknown', 'Soft Tissue Infections/Colonization', 'COVID-19', 'RTIs', 'Circulatory Conditions', 'Gastrointestinal Disorders', 'Unsorted Bacterial infections', 'Unknown', 'Unknown', 'Unknown', 'Unsorted Bacterial infections', 'Unknown', 'Bone Infection', 'Unsorted Bacterial infections', 'Unknown', 'Miscellaneous', 'Unknown', 'Porcine', 'RTIs', 'Unsorted Bacterial infections', 'Bone Infection', 'Unknown', 'Unsorted Bacterial infections', 'Soft Tissue Infections/Colonization', 'Unknown', 'UTIs', 'Unsorted Bacterial infections', 'Unknown', 'Head Injury and Traumatic Brain Injury', 'Unknown', 'Unknown', 'Head Injury and Traumatic Brain Injury', 'Soft Tissue Infections/Colonization', 'Unknown', 'Unknown', 'Unknown', 'RTIs', 'Unknown', 'Unknown', 'Unknown', 'Unsorted Bacterial infections', 'Unsorted Bacterial infections', 'Unknown', 'Unknown', 'Unsorted Bacterial infections', 'Unsorted Bacterial infections', 'RTIs', 'Unknown', 'Fruit', 'Unknown', 'Unsorted Bacterial infections', 'Unknown', 'Bone Infection', 'Unknown', 'Unknown', 'Unknown', 'Other Injuries', 'RTIs', 'Unknown', 'Unknown', 'RTIs', 'Miscellaneous', 'Bone Infection', 'Unknown', 'Unknown', 'Unsorted Bacterial infections', 'Immunodeficiency Disorders', 'Unsorted Bacterial infections', 'Soft Tissue Infections/Colonization', 'Unknown', 'RTIs', 'Other Injuries', 'Unknown', 'Bone Infection', 'Unsorted Bacterial infections', 'Soft Tissue Infections/Colonization', 'Unknown', 'RTIs', 'Unknown', 'Unsorted Bacterial infections', 'Unknown', 'RTIs', 'Unknown', 'Unknown', 'Unknown', 'Soft Tissue Infections/Colonization', 'Unknown', 'Unsorted Bacterial infections', 'Vascular Disorders', 'Unknown', 'Unknown', 'Unknown', 'Unknown', 'Unknown', 'Unknown', 'Unknown', 'Unknown', 'Unknown', 'Head Injury and Traumatic Brain Injury', 'Unknown', 'Unknown', 'Unknown', 'Bone Infection', 'Unsorted Bacterial infections', 'Soft Tissue Infections/Colonization', 'Unknown', 'Unknown', 'Unknown', 'Unsorted Bacterial infections', 'Unknown', 'Unknown', 'Bone Infection', 'Manured Lands', 'Unknown', 'Unsorted Bacterial infections', 'Unknown', 'Unknown', 'Unknown', 'Liver Diseases', 'Immunodeficiency Disorders', 'Immunodeficiency Disorders', 'Unknown', 'Unknown', 'Unknown', 'Unknown', 'Cardiac Disorders', 'Unknown', 'Unknown', 'Unknown', 'Unknown', 'Poultry', 'Unknown', 'Unknown', 'RTIs', 'Soft Tissue Infections/Colonization', 'RTIs', 'Unknown', 'Unknown', 'Unknown', 'Soft Tissue Infections/Colonization']
Source - manually curated: ['Solid Tumors', 'RTIs', 'Soft Tissue Infections/Colonization', 'Burns and Soft Tissue Injuries', 'RTIs', 'Soft Tissue Infections/Colonization', 'RTIs', 'Gastrointestinal Disorders', 'RTIs', 'Bacteremia', 'Gastrointestinal Disorders', 'RTIs', 'RTIs', 'Bacteremia', 'RTIs', 'RTIs', 'Unsorted Bacterial infections', 'RTIs', 'Gastrointestinal Disorders', 'Soft Tissue Infections/Colonization', 'Infusion and IV Equipment', 'RTIs', 'RTIs', 'Unknown', 'RTIs', 'RTIs', 'Soft Tissue Infections/Colonization', 'RTIs', 'RTIs', 'RTIs', 'Soft Tissue Infections/Colonization', 'RTIs', 'Unknown', 'RTIs', 'RTIs', 'Bacteremia', 'Bacteremia', 'RTIs', 'UTIs', 'Soft Tissue Infections/Colonization', 'Soft Tissue Infections/Colonization', 'Soft Tissue Infections/Colonization', 'RTIs', 'Others', 'Gastrointestinal Disorders', 'Bacteremia', 'Gastrointestinal Disorders', 'Soft Tissue Infections/Colonization', 'Soft Tissue Infections/Colonization', 'Soft Tissue Infections/Colonization', 'Unknown', 'Infusion and IV Equipment', 'RTIs', 'RTIs', 'Infusion and IV Equipment', 'Unknown', 'Others', 'RTIs', 'RTIs', 'RTIs', 'Burns and Soft Biopsy Injuries', 'Bacteremia', 'Unknown', 'UTIs', 'RTIs', 'UTIs', 'Soft Tissue Infections/Colonization', 'Burns and Soft Biopsy Injuries', 'Soft Tissue Infections/Colonization', 'RTIs', 'RTIs', 'Beds and Support Equipment', 'RTIs', 'Gastrointestinal Disorders', 'RTIs', 'Unsorted Bacterial infections', 'Influenza and other Respiratory Viral Infections', 'Unknown', 'RTIs', 'Soft Tissue Infections/Colonization', 'Gastrointestinal Disorders', 'RTIs', 'RTIs', 'RTIs', 'Soft Tissue Infections/Colonization', 'RTIs', 'Soft Tissue Infections/Colonization', 'Soft Tissue Infections/Colonization', 'RTIs', 'Pig', 'Miscellaneous', 'Surgical and Therapeutic Equipment', 'Bacteremia', 'Unsorted Bacterial infections', 'RTIs', 'UTIs', 'RTIs', 'Unsorted Bacterial infections', 'RTIs', 'RTIs', 'Cerebrovascular Diseases', 'Soft Tissue Infections/Colonization', 'Gastrointestinal Disorders', 'Cerebrovascular Diseases', 'UTIs', 'RTIs', 'Gastrointestinal Disorders', 'Soft Tissue Infections/Colonization', 'Bacteremia', 'Unsorted Bacterial infections', 'Gastrointestinal Disorders', 'Soft Tissue Infections/Colonization', 'RTIs', 'RTIs', 'RTIs', 'Gastrointestinal Disorders', 'Infusion and IV Equipment', 'Gastrointestinal Disorders', 'Bacteremia', 'RTIs', 'Miscellaneous', 'Soft Tissue Infections/Colonization', 'Soft Tissue Infections/Colonization', 'RTIs', 'Bacteremia', 'RTIs', 'Gastrointestinal Disorders', 'Soft Tissue Infections/Colonization', 'Soft Tissue Infections/Colonization', 'Bacteremia', 'Gastrointestinal Disorders', 'RTIs', 'Bacteremia', 'Gastrointestinal Disorders', 'Unknown', 'RTIs', 'RTIs', 'RTIs', 'RTIs', 'RTIs', 'UTIs', 'RTIs', 'Bacteremia', 'Soft Tissue Infections/Colonization', 'RTIs', 'Soft Tissue Infections/Colonization', 'Other Injuries', 'UTIs', 'Gastrointestinal Disorders', 'Bacteremia', 'RTIs', 'RTIs', 'Transplant Recepient', 'Bacteremia', 'Gastrointestinal Disorders', 'RTIs', 'RTIs', 'Gastrointestinal Disorders', 'Soft Tissue Infections/Colonization', 'Surgical and Therapeutic Equipment', 'RTIs', 'RTIs', 'Surgical and Therapeutic Equipment', 'RTIs', 'Soft Tissue Infections/Colonization', 'Soft Tissue Infections/Colonization', 'RTIs', 'RTIs', 'RTIs', 'Gastrointestinal Disorders', 'Cerebrovascular Diseases', 'RTIs', 'RTIs', 'RTIs', 'Unknown', 'RTIs', 'UTIs', 'RTIs', 'RTIs', 'Soft Tissue Infections/Colonization', 'RTIs', 'RTIs', 'RTIs', 'Bacteremia', 'Others', 'RTIs', 'Bacteremia', 'Soft Tissue Infections/Colonization', 'Gastrointestinal Disorders', 'RTIs', 'Solid Tumors', 'RTIs', 'RTIs', 'Beds and Support Equipment', 'Gastrointestinal Disorders', 'Soft Tissue Infections/Colonization', 'RTIs', 'Unknown', 'Gastrointestinal Disorders', 'Unsorted Bacterial infections', 'Soft Tissue Infections/Colonization', 'Soft Tissue Infections/Colonization', 'Bovine', 'RTIs', 'RTIs', 'Bacteremia', 'UTIs', 'Bacteremia', 'RTIs', 'RTIs', 'RTIs', 'Gastrointestinal Disorders']
Total entries: 2500
Total entries that didn't match when the metaminer curation was supported by database: 2
Total entries that didn't match when the metaminer curation was not supported by database: 212
{('homo sapiens', 'a. baumannii bacteremia (catheter-realated infection)', 'seoul national university bundang hospital intensive care unit'): ['Infusion and IV Equipment', 'Bacteremia'], ('homo sapiens', 'infection', 'trachy'): ['Unsorted Bacterial infections', 'Surgical and Therapeutic Equipment']}
{('homo sapiens', 'esophageal cancer', 'lavage fluid'): ['Gastrointestinal Disorders', 'Solid Tumors'], ('homo sapiens', None, 'trachea'): ['Unknown', 'RTIs'], ('homo sapiens', None, 'anal margin'): ['Unknown', 'Soft Tissue Infections/Colonization'], ('homo sapiens', 'burn', 'blood'): ['Bacteremia', 'Burns and Soft Tissue Injuries'], ('homo sapiens', 'mediastinal abscess', 'lavage fluid'): ['Gastrointestinal Disorders', 'RTIs'], ('homo sapiens', None, 'swab of axilla and groin'): ['Unknown', 'Soft Tissue Infections/Colonization'], ('homo sapiens', 'unknown', 'trach asp'): ['Unknown', 'RTIs'], ('homo sapiens', None, 'rectal'): ['Unknown', 'Gastrointestinal Disorders'], ('homo sapiens', None, 'tracheal aspirate'): ['Unknown', 'RTIs'], ('homo sapiens', 'acinetobacter baumannii infections', 'sputum'): ['RTIs', 'Bacteremia'], ('homo sapiens', 'missing', 'bronchial wash'): ['Allergic Disorders', 'RTIs'], ('homo sapiens', None, 'tracheal suction'): ['Unknown', 'RTIs'], ('homo sapiens', 'a. baumannii bacteremia (catheter-realated infection)', 'seoul national university bundang hospital intensive care unit'): ['Infusion and IV Equipment', 'Bacteremia'], ('homo sapiens', None, 'bronchial wash'): ['Allergic Disorders', 'RTIs'], ('homo sapiens', 'a. baumannii infection', 'periprosthetic liquid'): ['Unknown', 'Unsorted Bacterial infections'], ('homo sapiens', 'not applicable', 'c tra asp'): ['Unknown', 'RTIs'], ('homo sapiens', None, 'abdomen isolate'): ['Bone Infection', 'Gastrointestinal Disorders'], ('homo sapiens', None, 'swab_wound'): ['Other Injuries', 'Soft Tissue Infections/Colonization'], ('homo sapiens', 'urinary tract infection', 'urinary catheter'): ['UTIs', 'Infusion and IV Equipment'], ('homo sapiens', None, 'wash_bronchoalveolar lavage (bal)'): ['Immunodeficiency Disorders', 'RTIs'], ('homo sapiens', 'unknown', 'tracheal secretion'): ['Unknown', 'RTIs'], (None, None, 'respiratory'): ['RTIs', 'Unknown'], ('homo sapiens', 'bacterial infection', 'respiratory'): ['Unsorted Bacterial infections', 'RTIs'], ('homo sapiens', None, 'tracheal swab'): ['Unknown', 'RTIs'], ('homo sapiens', None, 'axilla/groin swab'): ['Unknown', 'Soft Tissue Infections/Colonization'], ('homo sapiens', 'unknown', 'endotracheal aspirate'): ['Unknown', 'RTIs'], ('homo sapiens', None, 'tracheal aspirate isolate'): ['Unknown', 'RTIs'], ('homo sapiens', None, 'swab (wound)'): ['Other Injuries', 'Soft Tissue Infections/Colonization'], ('homo sapiens', 'unknown', 'tracheal aspirate'): ['Unknown', 'RTIs'], ('homo sapiens', 'pneumonia', 'hospital environment'): ['RTIs', 'Unknown'], ('homo sapiens', 'not provided', 'tracheal aspirate'): ['Unknown', 'RTIs'], ('homo sapiens', 'missing', 'tracheal aspirate'): ['Unknown', 'RTIs'], ('homo sapiens', 'not applicable', 'lung'): ['Unknown', 'RTIs'], ('homo sapiens', None, 'catheter urine'): ['Infusion and IV Equipment', 'UTIs'], ('homo sapiens', None, 'axilla and groin swab'): ['Unknown', 'Soft Tissue Infections/Colonization'], ('homo sapiens', None, 'wound isolate'): ['Bone Infection', 'Soft Tissue Infections/Colonization'], ('homo sapiens', None, 'endotracheal aspirate'): ['Unknown', 'RTIs'], ('not applicable', 'not applicable', 'background grass (no manure on field)'): ['Agricultural Soil', 'Others'], ('homo sapiens', 'colonisation', 'rectal'): ['Unsorted Bacterial infections', 'Gastrointestinal Disorders'], ('homo sapiens', 'acinetobacter baumannii infections', 'broncho-alveolar lavage'): ['RTIs', 'Bacteremia'], ('homo sapiens', None, 'skin; arms/legs'): ['Unknown', 'Soft Tissue Infections/Colonization'], ('homo sapiens', None, 'axilla/groin'): ['Unknown', 'Soft Tissue Infections/Colonization'], ('homo sapiens', 'bloodstream infection', 'foley catheter'): ['Bacteremia', 'Infusion and IV Equipment'], ('homo sapiens', 'unknown', 'central venous access devices'): ['Unknown', 'Infusion and IV Equipment'], ('homo sapiens', 'burnt patient', 'uroculture'): ['Soft Tissue Infections/Colonization', 'Burns and Soft Biopsy Injuries'], ('homo sapiens', 'osteomyelitis', 'bone'): ['Bone Infection', 'Unknown'], ('homo sapiens', 'surgical injuries', 'urine'): ['Soft Tissue Infections/Colonization', 'UTIs'], ('homo sapiens', 'not applicable', 'trach asp'): ['Unknown', 'RTIs'], ('homo sapiens', 'burn infection', 'burn patient'): ['CNS infections', 'Burns and Soft Biopsy Injuries'], ('homo sapiens', 'vap/hap', 'tracheal aspirate'): ['Unknown', 'RTIs'], (None, None, 'bedside rail in hospital intensive care unit'): ['Miscellaneous', 'Beds and Support Equipment'], ('homo sapiens', None, 'abdominal wound isolate'): ['Soft Tissue Infections/Colonization', 'Gastrointestinal Disorders'], ('homo sapiens', 'covid-19, acinetobacter pneumonia', 'sputum'): ['COVID-19', 'RTIs'], ('homo sapiens', 'infection', 'hydrothorax'): ['RTIs', 'Unsorted Bacterial infections'], ('homo sapiens', 'h1n1', 'sputum'): ['Circulatory Conditions', 'Influenza and other Respiratory Viral Infections'], ('homo sapiens', None, 'secreta'): ['Gastrointestinal Disorders', 'Unknown'], ('homo sapiens', 'infection', 'etta'): ['Unsorted Bacterial infections', 'RTIs'], ('homo sapiens', 'not determined', 'penis swab'): ['Unknown', 'Soft Tissue Infections/Colonization'], ('homo sapiens', 'acinetobacter infections', 'sput'): ['Unsorted Bacterial infections', 'RTIs'], ('homo sapiens', 'infection', 'tracheal secretion'): ['Unsorted Bacterial infections', 'RTIs'], ('homo sapiens', None, 'right shin blister'): ['Unknown', 'Soft Tissue Infections/Colonization'], ('homo sapiens', 'nicu', 'hand'): ['Miscellaneous', 'Soft Tissue Infections/Colonization'], ('sus scrofa domesticus', 'missing', 'swab'): ['Porcine', 'Pig'], ('homo sapiens', 'pneumonia', 'drainage basin water'): ['RTIs', 'Miscellaneous'], ('homo sapiens', 'infection', 'trachy'): ['Unsorted Bacterial infections', 'Surgical and Therapeutic Equipment'], ('homo sapiens', None, 'blood isolate'): ['Bone Infection', 'Bacteremia'], ('homo sapiens', 'not applicable', 'oral swab_p101_bu22'): ['Unknown', 'Unsorted Bacterial infections'], ('homo sapiens', 'missing', 'urethral swab'): ['UTIs', 'Unsorted Bacterial infections'], ('homo sapiens', None, 'low respiratory tract'): ['Unknown', 'RTIs'], ('homo sapiens', 'cerebral infarction', 'sputum'): ['Head Injury and Traumatic Brain Injury', 'Cerebrovascular Diseases'], ('homo sapiens', None, 'foot'): ['Unknown', 'Soft Tissue Infections/Colonization'], ('homo sapiens', 'not applicable', 'oral swab_p23_bu25'): ['Unknown', 'Unsorted Bacterial infections'], ('homo sapiens', None, 'heel'): ['Unknown', 'Soft Tissue Infections/Colonization'], ('homo sapiens', 'infection', 'respiratory'): ['Unsorted Bacterial infections', 'RTIs'], ('homo sapiens', None, 'trachael aspirate'): ['Unknown', 'RTIs'], ('homo sapiens', 'infection', 'tip of intravascular catheter'): ['Unsorted Bacterial infections', 'Infusion and IV Equipment'], ('homo sapiens', 'severe pneumonia, bloodstream infection, bone marrow infection, septic shock', 'blood'): ['RTIs', 'Bacteremia'], (None, None, 'coconut meat [foodon:00003856]; ready-to-eat (rte) [foodon:03316636]; food (chunks) [foodon:00004555]; food (frozen) [foodon:03302148]; retail environment [envo:01001448]'): ['Fruit', 'Miscellaneous'], ('homo sapiens', 'acinetobacter baumannii skin abscess infection', 'missing'): ['Unknown', 'Soft Tissue Infections/Colonization'], ('homo sapiens', 'infection', 'hip'): ['Unsorted Bacterial infections', 'Soft Tissue Infections/Colonization'], ('homo sapiens', None, 'tracheal secretion'): ['Unknown', 'RTIs'], ('homo sapiens', 'surveillance', 'rectal swab'): ['Miscellaneous', 'Gastrointestinal Disorders'], ('homo sapiens', None, 'tissue;sacrum'): ['Bone Infection', 'Unknown'], ('homo sapiens', 'vap', 'ett'): ['Unknown', 'RTIs'], ('homo sapiens', None, 'tracheal aspirate/wash'): ['Immunodeficiency Disorders', 'RTIs'], ('homo sapiens', 'infection', 'severe trauma site'): ['Unsorted Bacterial infections', 'Other Injuries'], ('homo sapiens', None, 'draining liquid of kidney transplantation recipient'): ['Unknown', 'Transplant Recepient'], ('homo sapiens', 'not applicable', 'rectal swab_p78_bu13'): ['Unknown', 'Gastrointestinal Disorders'], ('homo sapiens', 'not applicable', 'bronch wash'): ['Unknown', 'RTIs'], ('homo sapiens', None, 'perirectal'): ['Soft Tissue Infections/Colonization', 'Gastrointestinal Disorders'], ('homo sapiens', 'acinetobacter infection', 'endobronchial tube'): ['Unsorted Bacterial infections', 'Surgical and Therapeutic Equipment'], ('homo sapiens', 'cerebrovascular disease', 'sputum'): ['Vascular Disorders', 'RTIs'], ('homo sapiens', 'endotracheal infection', 'endotracheal tube'): ['Unknown', 'Surgical and Therapeutic Equipment'], ('homo sapiens', 'missing', 'left hip'): ['Unknown', 'Soft Tissue Infections/Colonization'], ('homo sapiens', None, 'abdomen'): ['Unknown', 'Gastrointestinal Disorders'], ('homo sapiens', 'not determined', 'tracheal secretions'): ['Unknown', 'RTIs'], ('homo sapiens', None, 'other isolate'): ['Bone Infection', 'Unknown'], ('homo sapiens', 'a. baumannii infection', 'endotracheal aspirate'): ['Unknown', 'RTIs'], ('homo sapiens', None, 'leg, left'): ['Unknown', 'Soft Tissue Infections/Colonization'], ('homo sapiens', None, 'tissue_lung'): ['Unknown', 'RTIs'], ('not applicable', 'not applicable', 'control grass (no manure)'): ['Manured Lands', 'Others'], ('homo sapiens', 'bilateral lung contusion and infection,bloodstream infection,abdominal infection with peritonitis,urinary tract infection', 'skin damage'): ['Unsorted Bacterial infections', 'Bacteremia'], ('homo sapiens', None, 'skin; arms and legs'): ['Unknown', 'Soft Tissue Infections/Colonization'], ('homo sapiens', None, 'lung'): ['Unknown', 'RTIs'], ('homo sapiens', 'liver metastases', 'pleural effusion'): ['Liver Diseases', 'Solid Tumors'], (None, None, 'crash trolley 2_na_room 7'): ['Unknown', 'Beds and Support Equipment'], ('homo sapiens', None, 'scrotal'): ['Unknown', 'Soft Tissue Infections/Colonization'], ('homo sapiens', 'not collected', 'st'): ['Cardiac Disorders', 'Unknown'], ('homo sapiens', 'not applicable', 'rectal swab_p26_bu28'): ['Unknown', 'Gastrointestinal Disorders'], ('homo sapiens', 'not applicable', 'oral swab_p96_bu15'): ['Unknown', 'Unsorted Bacterial infections'], ('turkey', None, None): ['Poultry', 'Bovine'], ('homo sapiens', None, 'chest'): ['Unknown', 'RTIs'], ('homo sapiens', 'aecopd', 'blood'): ['RTIs', 'Bacteremia'], ('homo sapiens', 'severe pneumonia,septic shock', 'sputamentum'): ['RTIs', 'Bacteremia'], ('homo sapiens', None, 'throat wash'): ['Unknown', 'RTIs'], ('homo sapiens', None, 'perirectal swab'): ['Soft Tissue Infections/Colonization', 'Gastrointestinal Disorders']}
```

In [24]:

```
# next, the sample! 

sample_entry = []
sample_entry_wo_d = []
sample_metaminer = []
sample_metaminer_wo_d = []
sample_manual = []
sample_manual_wo_d = []

number = {}
for i in range(len(normalized_cleaned_data)):
    if normalized_cleaned_data.loc[i,'sample'] == normalized_cleaned_data.loc[i,'Manual_sample']:
        pass
    else:
        sample_entry.append(normalized_cleaned_data.loc[i,'isolation_source'])
        sample_metaminer.append(normalized_cleaned_data.loc[i,'sample'])
        sample_manual.append(normalized_cleaned_data.loc[i,'Manual_sample'])
        key = (normalized_cleaned_data.loc[i, 'host'],
               normalized_cleaned_data.loc[i, 'host_disease'],
               normalized_cleaned_data.loc[i, 'isolation_source'])

        value = [normalized_cleaned_data.loc[i, 'sample'], 
                 normalized_cleaned_data.loc[i, 'Manual_sample']]
        
        number[key] = value

number1 = {}
for i in range(len(normalized_cleaned_data)):
    if normalized_cleaned_data.loc[i,'sample_d'] == normalized_cleaned_data.loc[i,'Manual_sample']:
        pass
    else:
        sample_entry_wo_d.append(normalized_cleaned_data.loc[i,'isolation_source'])
        sample_metaminer_wo_d.append(normalized_cleaned_data.loc[i,'sample_d'])
        sample_manual_wo_d.append(normalized_cleaned_data.loc[i,'Manual_sample'])
        key = (normalized_cleaned_data.loc[i, 'host'],
               normalized_cleaned_data.loc[i, 'host_disease'],
               normalized_cleaned_data.loc[i, 'isolation_source'])

        value = [normalized_cleaned_data.loc[i, 'sample_d'], 
                 normalized_cleaned_data.loc[i, 'Manual_sample']]
        
        number1[key] = value


print(f"Source submitted entry: {sample_entry}")
print(f"Source - metaminer curated with database: {sample_metaminer}")
print(f"Source - manually curated: {sample_manual}")

print(f"Source submitted entry without database: {sample_entry_wo_d}")
print(f"Source - metaminer curated without database: {sample_metaminer_wo_d}")
print(f"Source - manually curated: {sample_manual_wo_d}")

total_entries = len(pd.notnull(normalized_cleaned_data['isolation_source']))
Total_didnt_match = len(sample_metaminer)
Total_didnt_match_wo_d = len(sample_metaminer_wo_d)

print(f"Total entries: {total_entries}")
print(f"Total entries that didn't match when the metaminer curation was supported by database: {Total_didnt_match}")
print(f"Total entries that didn't match when the metaminer curation was not supported by database: {Total_didnt_match_wo_d}")

print(number)
print(number1)
```

```
Source submitted entry: ['seoul national university bundang hospital intensive care unit', 'trachy']
Source - metaminer curated with database: ['Catheter', 'Unknown']
Source - manually curated: ['Unknown', 'Tracheostomy Tube']
Source submitted entry without database: ['lavage fluid', 'trachea', 'anal margin', 'lavage fluid', 'alcohol foam dispenser in hospital intensive care unit', 'urinary tract', 'swab of axilla and groin', 'drain fluid', 'rectal', 'hospital', 'rectal', 'missing', 'patient in hospital', 'seoul national university bundang hospital intensive care unit', 'diabetic foot', 'oral swab', 'periprosthetic liquid', 'c tra asp', 'abscess; hip right', 'urine', 'swab_wound', 'urinary catheter', 'blood', 'axilla/groin swab', 'urine', 'patient sputum', 'tracheal aspirate isolate', 'blood specimen', 'swab (wound)', 'abscess tissue', 'blood', 'lung', 'catheter urine', 'axilla/groin swab', 'surfaces surveillance', 'axilla and groin swab', 'bronchoaspirate', 'background grass (no manure on field)', 'rectal', 'rectal', 'skin/wound', 'skin; arms/legs', 'blood', 'axilla/groin', 'foley catheter', 'central venous access devices', 'background grass (no manure on field)', 'bloodstream', 'lung', 'tissues', 'skin and soft tissue from inpatient', 'uroculture', 'bone', 'blood', 'urine', 'urine', 'axilla and groin swab', 'swab of axilla and groin', 'axilla and groin swab', 'blood', 'bedside rail in hospital intensive care unit', 'bronchoalveolar wash', 'secreta', 'penis swab', 'rectal', 'skin/wound', 'sput', 'right shin blister', 'hand', 'sputum', 'soft tissue', 'drainage basin water', 'trachy', 'oral swab_p101_bu22', 'wound', 'urine', 'axilla and groin swab', 'rectal', 'urine', 'rectal', 'foot', 'oral swab_p23_bu25', 'rectal', 'heel', 'rectal', 'joint', 'tip of intravascular catheter', 'rectal', 'blood', 'unknown', 'coconut meat [foodon:00003856]; ready-to-eat (rte) [foodon:03316636]; food (chunks) [foodon:00004555]; food (frozen) [foodon:03302148]; retail environment [envo:01001448]', 'missing', 'right leg muscle, right leg fascia, central venous catheter, dialysis catheter, surface of the left leg ulcer, and left lower leg', 'rectal', 'axilla/groin swab', 'swab_wound', 'rectal', 'blood', 'rectal swab', 'trachea', 'blood', 'urine', 'blood', 'swab (wound)', 'blood', 'tracheal aspirate isolate', 'urine', 'rectal', 'alcohol foam dispenser in hospital intensive care unit', 'rectal swab_p78_bu13', 'bronch wash', 'axilla/groin swab', 'endobronchial tube', 'endotracheal tube', 'axilla and groin swab', 'left hip', 'urine', 'urine', 'endotracheal throat swab', 'leg, left', 'surfaces surveillance', 'sputum specimen', 'control grass (no manure)', 'trachea', 'skin damage', 'sputum', 'skin; arms and legs', 'endovascular', 'rectal', 'lung', 'skin/wound', 'lung', 'crash trolley 2_na_room 7', 'rectal', 'blood', 'scrotal', 'endotracheal throat swab', 'rectal swab_p26_bu28', 'oral swab_p96_bu15', 'axilla/groin swab', 'axilla and groin swab', 'skin/wound', 'urine', 'blood', 'throat wash']
Source - metaminer curated without database: ['Unknown', 'Unknown', 'Unknown', 'Pus', 'Disinfectant', 'Unknown', 'Swab', 'Wound Swab', 'Unknown', 'Blood', 'Unknown', 'Skin', 'Blood', 'Catheter', 'Unknown', 'Swab', 'Unknown', 'Unknown', 'Pus', 'Hair', 'Swab', 'Unknown', 'Blood', 'Swab', 'Hair', 'Sputum', 'Unknown', 'Blood', 'Swab', 'Biopsy', 'Blood', 'Unknown', 'Catheter', 'Swab', 'Unknown', 'Swab', 'Unknown', 'Agricultural Soil', 'Unknown', 'Unknown', 'Skin', 'Skin', 'Blood', 'Unknown', 'Unknown', 'Unknown', 'Agricultural Soil', 'Unknown', 'Unknown', 'Wound Swab', 'Skin', 'Unknown', 'Bone', 'Blood', 'Wound Swab', 'Wound Swab', 'Swab', 'Swab', 'Swab', 'Blood', 'Hospital surface', 'Unknown', 'Stool', 'Swab', 'Unknown', 'Skin', 'Unknown', 'Unknown', 'Hospital surface', 'Sputum', 'Fat Tissue', 'Amniotic Fluid', 'Unknown', 'Swab', 'Fat Tissue', 'Wound Swab', 'Swab', 'Unknown', 'Wound Swab', 'Unknown', 'Unknown', 'Swab', 'Unknown', 'Unknown', 'Unknown', 'Unknown', 'Unknown', 'Unknown', 'Bone', 'Skin', 'Fruit', 'Skin', 'Muscle', 'Unknown', 'Swab', 'Swab', 'Unknown', 'Blood', 'Hospital surface', 'Unknown', 'Blood', 'Wound Swab', 'Blood', 'Swab', 'Blood', 'Unknown', 'Wound Swab', 'Unknown', 'Disinfectant', 'Swab', 'Unknown', 'Swab', 'Unknown', 'Tracheal', 'Swab', 'Unknown', 'Hair', 'Wound Swab', 'Swab', 'Unknown', 'Unknown', 'Sputum', 'Manured Lands', 'Unknown', 'Skin', 'Sputum', 'Skin', 'Unknown', 'Unknown', 'Unknown', 'Skin', 'Unknown', 'Unknown', 'Unknown', 'Blood', 'Unknown', 'Swab', 'Swab', 'Swab', 'Swab', 'Swab', 'Skin', 'Wound Swab', 'Blood', 'Unknown']
Source - manually curated: ['Bronchoalveolar Lavage', 'Tracheal', 'Perianal Swab', 'Bronchoalveolar Lavage', 'Disinfectant Dispenser', 'Urine', 'Skin Swab', 'Unknown', 'Rectal Swab', 'Unknown', 'Rectal Swab', 'Unknown', 'Unknown', 'Unknown', 'Biopsy', 'Unknown', 'Biopsy', 'Tracheal', 'Wound Swab', 'Urine', 'Wound Swab', 'Catheter', 'Stool', 'Axillary Swab', 'Urine', 'Unknown', 'Tracheal', 'Unknown', 'Wound Swab', 'Wound Swab', 'Unknown', 'Biopsy', 'Urine', 'Axillary Swab', 'Swab', 'Skin Swab', 'Bronchoalveolar Lavage', 'Others', 'Rectal Swab', 'Rectal Swab', 'Wound Swab', 'Skin Swab', 'Unknown', 'Skin Swab', 'Catheter', 'Catheter', 'Others', 'Blood', 'Biopsy', 'Biopsy', 'Biopsy', 'Urine', 'Unknown', 'Unknown', 'Urine', 'Urine', 'Skin Swab', 'Skin Swab', 'Skin Swab', 'Unknown', 'Bed', 'Bronchoalveolar Lavage', 'Unknown', 'Skin Swab', 'Rectal Swab', 'Wound Swab', 'Sputum', 'Wound Swab', 'Skin Swab', 'Tracheal', 'Biopsy', 'Hospital surface', 'Tracheostomy Tube', 'Oral Swab', 'Wound Swab', 'Urine', 'Skin Swab', 'Rectal Swab', 'Urine', 'Rectal Swab', 'Skin Swab', 'Oral Swab', 'Rectal Swab', 'Skin Swab', 'Rectal Swab', 'Synovial Fluid', 'Catheter', 'Rectal Swab', 'Blood', 'Skin Swab', 'Miscellaneous', 'Pus', 'Wound Swab', 'Rectal Swab', 'Axillary Swab', 'Wound Swab', 'Rectal Swab', 'Unknown', 'Rectal Swab', 'Tracheal', 'Unknown', 'Urine', 'Unknown', 'Wound Swab', 'Unknown', 'Tracheal', 'Urine', 'Rectal Swab', 'Disinfectant Dispenser', 'Rectal Swab', 'Bronchoalveolar Lavage', 'Axillary Swab', 'Endobronchial Tube', 'Endotracheal Tube', 'Skin Swab', 'Skin Swab', 'Urine', 'Urine', 'Throat Swab', 'Skin Swab', 'Swab', 'Unknown', 'Others', 'Tracheal', 'Wound Swab', 'Tracheal', 'Skin Swab', 'Blood', 'Rectal Swab', 'Biopsy', 'Wound Swab', 'Biopsy', 'Care Cart', 'Rectal Swab', 'Unknown', 'Skin Swab', 'Throat Swab', 'Rectal Swab', 'Oral Swab', 'Axillary Swab', 'Skin Swab', 'Wound Swab', 'Urine', 'Unknown', 'Throat Swab']
Total entries: 2500
Total entries that didn't match when the metaminer curation was supported by database: 2
Total entries that didn't match when the metaminer curation was not supported by database: 145
{('homo sapiens', 'a. baumannii bacteremia (catheter-realated infection)', 'seoul national university bundang hospital intensive care unit'): ['Catheter', 'Unknown'], ('homo sapiens', 'infection', 'trachy'): ['Unknown', 'Tracheostomy Tube']}
{('homo sapiens', 'esophageal cancer', 'lavage fluid'): ['Unknown', 'Bronchoalveolar Lavage'], ('homo sapiens', None, 'trachea'): ['Unknown', 'Tracheal'], ('homo sapiens', None, 'anal margin'): ['Unknown', 'Perianal Swab'], ('homo sapiens', 'mediastinal abscess', 'lavage fluid'): ['Pus', 'Bronchoalveolar Lavage'], (None, None, 'alcohol foam dispenser in hospital intensive care unit'): ['Disinfectant', 'Disinfectant Dispenser'], ('homo sapiens', 'urinary tract infection', 'urinary tract'): ['Unknown', 'Urine'], ('homo sapiens', None, 'swab of axilla and groin'): ['Swab', 'Skin Swab'], ('homo sapiens', 'peptic ulcer, gastric perforation', 'drain fluid'): ['Wound Swab', 'Unknown'], ('homo sapiens', None, 'rectal'): ['Unknown', 'Rectal Swab'], ('homo sapiens', 'blood stream infection', 'hospital'): ['Blood', 'Unknown'], ('homo sapiens', 'skin and soft tissue infection', 'missing'): ['Skin', 'Unknown'], ('homo sapiens', 'blood stream infection', 'patient in hospital'): ['Blood', 'Unknown'], ('homo sapiens', 'a. baumannii bacteremia (catheter-realated infection)', 'seoul national university bundang hospital intensive care unit'): ['Catheter', 'Unknown'], ('homo sapiens', 'diabetic foot', 'diabetic foot'): ['Unknown', 'Biopsy'], (None, None, 'oral swab'): ['Swab', 'Unknown'], ('homo sapiens', 'a. baumannii infection', 'periprosthetic liquid'): ['Unknown', 'Biopsy'], ('homo sapiens', 'not applicable', 'c tra asp'): ['Unknown', 'Tracheal'], ('homo sapiens', None, 'abscess; hip right'): ['Pus', 'Wound Swab'], ('cat', 'uti', 'urine'): ['Hair', 'Urine'], ('homo sapiens', None, 'swab_wound'): ['Swab', 'Wound Swab'], ('homo sapiens', 'urinary tract infection', 'urinary catheter'): ['Unknown', 'Catheter'], ('homo sapiens', 'not available', 'blood'): ['Blood', 'Stool'], ('homo sapiens', None, 'axilla/groin swab'): ['Swab', 'Axillary Swab'], ('cat', None, 'urine'): ['Hair', 'Urine'], ('homo sapiens', 'pneumonia', 'patient sputum'): ['Sputum', 'Unknown'], ('homo sapiens', None, 'tracheal aspirate isolate'): ['Unknown', 'Tracheal'], ('homo sapiens', 'hospital acquired infection', 'blood specimen'): ['Blood', 'Unknown'], ('homo sapiens', None, 'swab (wound)'): ['Swab', 'Wound Swab'], ('homo sapiens', 'hopsital aqcuired infection', 'abscess tissue'): ['Biopsy', 'Wound Swab'], ('homo sapiens', 'missing', 'blood'): ['Blood', 'Unknown'], ('homo sapiens', 'not applicable', 'lung'): ['Unknown', 'Biopsy'], ('homo sapiens', None, 'catheter urine'): ['Catheter', 'Urine'], ('homo sapiens', 'colonization', 'surfaces surveillance'): ['Unknown', 'Swab'], ('homo sapiens', None, 'axilla and groin swab'): ['Swab', 'Skin Swab'], ('homo sapiens', 'nosocomial infection', 'bronchoaspirate'): ['Unknown', 'Bronchoalveolar Lavage'], ('not applicable', 'not applicable', 'background grass (no manure on field)'): ['Agricultural Soil', 'Others'], ('homo sapiens', 'colonisation', 'rectal'): ['Unknown', 'Rectal Swab'], ('homo sapiens', None, 'skin/wound'): ['Skin', 'Wound Swab'], ('homo sapiens', None, 'skin; arms/legs'): ['Skin', 'Skin Swab'], ('homo sapiens', None, 'axilla/groin'): ['Unknown', 'Skin Swab'], ('homo sapiens', 'bloodstream infection', 'foley catheter'): ['Unknown', 'Catheter'], ('homo sapiens', 'unknown', 'central venous access devices'): ['Unknown', 'Catheter'], ('homo sapiens', 'acute myeloid leukemia m2', 'bloodstream'): ['Unknown', 'Blood'], ('homo sapiens', 'wound infection', 'tissues'): ['Wound Swab', 'Biopsy'], ('homo sapiens', 'skin and soft tissue', 'skin and soft tissue from inpatient'): ['Skin', 'Biopsy'], ('homo sapiens', 'burnt patient', 'uroculture'): ['Unknown', 'Urine'], ('homo sapiens', 'osteomyelitis', 'bone'): ['Bone', 'Unknown'], ('homo sapiens', 'surgical injuries', 'urine'): ['Wound Swab', 'Urine'], ('homo sapiens', 'hopsital aqcuired infection', 'swab of axilla and groin'): ['Swab', 'Skin Swab'], (None, None, 'bedside rail in hospital intensive care unit'): ['Hospital surface', 'Bed'], ('homo sapiens', 'pneumonia', 'bronchoalveolar wash'): ['Unknown', 'Bronchoalveolar Lavage'], ('homo sapiens', None, 'secreta'): ['Stool', 'Unknown'], ('homo sapiens', 'not determined', 'penis swab'): ['Swab', 'Skin Swab'], ('homo sapiens', 'acinetobacter infections', 'sput'): ['Unknown', 'Sputum'], ('homo sapiens', None, 'right shin blister'): ['Unknown', 'Wound Swab'], ('homo sapiens', 'nicu', 'hand'): ['Hospital surface', 'Skin Swab'], ('homo sapiens', 'respiratory failure', 'sputum'): ['Sputum', 'Tracheal'], ('homo sapiens', None, 'soft tissue'): ['Fat Tissue', 'Biopsy'], ('homo sapiens', 'pneumonia', 'drainage basin water'): ['Amniotic Fluid', 'Hospital surface'], ('homo sapiens', 'infection', 'trachy'): ['Unknown', 'Tracheostomy Tube'], ('homo sapiens', 'not applicable', 'oral swab_p101_bu22'): ['Swab', 'Oral Swab'], ('homo sapiens', 'soft tissue', 'wound'): ['Fat Tissue', 'Wound Swab'], ('homo sapiens', None, 'foot'): ['Unknown', 'Skin Swab'], ('homo sapiens', 'not applicable', 'oral swab_p23_bu25'): ['Swab', 'Oral Swab'], ('homo sapiens', None, 'heel'): ['Unknown', 'Skin Swab'], ('homo sapiens', None, 'joint'): ['Unknown', 'Synovial Fluid'], ('homo sapiens', 'infection', 'tip of intravascular catheter'): ['Unknown', 'Catheter'], ('homo sapiens', 'severe pneumonia, bloodstream infection, bone marrow infection, septic shock', 'blood'): ['Bone', 'Blood'], ('homo sapiens', 'skin infection', 'unknown'): ['Skin', 'Skin Swab'], (None, None, 'coconut meat [foodon:00003856]; ready-to-eat (rte) [foodon:03316636]; food (chunks) [foodon:00004555]; food (frozen) [foodon:03302148]; retail environment [envo:01001448]'): ['Fruit', 'Miscellaneous'], ('homo sapiens', 'acinetobacter baumannii skin abscess infection', 'missing'): ['Skin', 'Pus'], ('homo sapiens', 'necrotizing fasciitis', 'right leg muscle, right leg fascia, central venous catheter, dialysis catheter, surface of the left leg ulcer, and left lower leg'): ['Muscle', 'Wound Swab'], ('homo sapiens', 'surveillance', 'rectal swab'): ['Hospital surface', 'Rectal Swab'], ('homo sapiens', 'not applicable', 'rectal swab_p78_bu13'): ['Swab', 'Rectal Swab'], ('homo sapiens', 'not applicable', 'bronch wash'): ['Unknown', 'Bronchoalveolar Lavage'], ('homo sapiens', 'acinetobacter infection', 'endobronchial tube'): ['Unknown', 'Endobronchial Tube'], ('homo sapiens', 'endotracheal infection', 'endotracheal tube'): ['Tracheal', 'Endotracheal Tube'], ('homo sapiens', 'missing', 'left hip'): ['Unknown', 'Skin Swab'], ('homo sapiens', 'infections,nosocomial', 'endotracheal throat swab'): ['Swab', 'Throat Swab'], ('homo sapiens', None, 'leg, left'): ['Unknown', 'Skin Swab'], ('homo sapiens', 'hospital acquired infection', 'sputum specimen'): ['Sputum', 'Unknown'], ('not applicable', 'not applicable', 'control grass (no manure)'): ['Manured Lands', 'Others'], ('homo sapiens', 'bilateral lung contusion and infection,bloodstream infection,abdominal infection with peritonitis,urinary tract infection', 'skin damage'): ['Skin', 'Wound Swab'], ('homo sapiens', None, 'skin; arms and legs'): ['Skin', 'Skin Swab'], ('homo sapiens', 'bacteremia', 'endovascular'): ['Unknown', 'Blood'], ('homo sapiens', None, 'lung'): ['Unknown', 'Biopsy'], ('homo sapiens', 'pneumonia', 'lung'): ['Unknown', 'Biopsy'], (None, None, 'crash trolley 2_na_room 7'): ['Unknown', 'Care Cart'], ('homo sapiens', None, 'scrotal'): ['Unknown', 'Skin Swab'], ('homo sapiens', 'infections, nosocomial', 'endotracheal throat swab'): ['Swab', 'Throat Swab'], ('homo sapiens', 'not applicable', 'rectal swab_p26_bu28'): ['Swab', 'Rectal Swab'], ('homo sapiens', 'not applicable', 'oral swab_p96_bu15'): ['Swab', 'Oral Swab'], ('homo sapiens', None, 'throat wash'): ['Unknown', 'Throat Swab']}
```

In [25]:

```
match_df = pd.DataFrame({
    'Category': ['Identified Host', 'Source Category', 'Source', 'Sample'],
    'Matched (With DB)': [2500, 2498, 2498, 2498],
    'Matched (Without DB)': [2494, 2475, 2288, 2355]
})

matrix = match_df.set_index('Category').T 

fig = px.imshow(matrix, 
        labels=dict(x="Category", y="Method", color="Count"),
        x=matrix.columns, y=matrix.index, 
        color_continuous_scale='YlGnBu',
        text_auto=True)

fig.update_layout(
    font=dict(size=14, color='black'), 
    xaxis=dict(
        title="Category Level",
        titlefont=dict(size=16, color='black'),
        tickfont=dict(size=14, color='black'),
        showline=True,
        linecolor='black',
        ticks='outside',
        tickangle = -45,
        tickwidth=2,
        ticklen=6,
    ),
    yaxis=dict(
        title = None,
        tickfont=dict(size=14, color='black'),
        showline=True,
        linecolor='black',
        ticks='outside',
        tickwidth=2,
        ticklen=6,
    ),
    height=450,
    width=500,
    showlegend=False,  
    plot_bgcolor="rgba(0, 0, 0, 0)",
    paper_bgcolor="rgba(0, 0, 0, 0)",
    coloraxis_showscale=False
)

# fig.show()

fig.write_image("./metaminer_bioarchiv/abau_comparison/isolation_source_normalized_2.png", format="png", scale=4)
```

- Sequencing Technologies used for sequencing:

In [26]:

```
# extracting just the raw values, metaminer normalized values and Manually curated values
validate_sequencing_technology_df = normalized_cleaned_data[['Sequencing_Technology', 'Categorized_sequencing_technologies', 'manually_curated_sequencing_technology']]
# print(validate_sequencing_technology_df[validate_sequencing_technology_df['Categorized_sequencing_technologies'] == 'Ion Torrent'])

normalization_didt_match = []
metaminer_standardized = []
manually_standardized = []

for i in range(len(validate_sequencing_technology_df)):
    if validate_sequencing_technology_df.iloc[i, 1] == validate_sequencing_technology_df.iloc[i, 2]:
        continue
    else:
        normalization_didt_match.append(validate_sequencing_technology_df.iloc[i,0])
        metaminer_standardized.append(validate_sequencing_technology_df.iloc[i, 1])
        manually_standardized.append(validate_sequencing_technology_df.iloc[i, 2])

print(f"Not matched entries: {normalization_didt_match}")
print(f"Metaminer curated entries for non-matched values: {metaminer_standardized}")
print(f"Manual curated entries for non-matched values: {manually_standardized}")

total_entries = len(pd.notnull(validate_sequencing_technology_df['Sequencing_Technology']))
Total_didnt_match = len(normalization_didt_match)

print(f"Total entries: {total_entries}")
print(f"Total entries that didn't match: {Total_didnt_match}")
```

```
Not matched entries: ['GenoLab M', 'Genelab M', 'Genelab M', 'Genelab M']
Metaminer curated entries for non-matched values: ['Unknown', 'Unknown', 'Unknown', 'Unknown']
Manual curated entries for non-matched values: ['GenoLab M', 'GenoLab M', 'GenoLab M', 'GenoLab M']
Total entries: 2500
Total entries that didn't match: 4
```

In [27]:

```
# plotting 

match_df = pd.DataFrame({
    'Category': ['Sequencing Technologies'],
    'Matched': [2496],
})

matrix = match_df.set_index('Category').T 

fig = px.imshow(matrix, 
        labels=dict(x="Category", y="Method", color="Count"),
        x=matrix.columns, y=matrix.index, 
        color_continuous_scale='Aggrnyl',
        text_auto=True)

fig.update_layout(
    font=dict(size=14, color='black'), 
    xaxis=dict(
        title=None,
        titlefont=dict(size=16, color='black'),
        tickfont=dict(size=14, color='black'),
        showline=True,
        linecolor='black',
        ticks='outside',
        # tickangle = -45,
        tickwidth=2,
        ticklen=6,
    ),
    yaxis=dict(
        title = None,
        tickfont=dict(size=14, color='black'),
        showline=True,
        linecolor='black',
        ticks='outside',
        tickwidth=2,
        ticklen=6,
    ),
    height=450,
    width=500,
    showlegend=False,  
    plot_bgcolor="rgba(0, 0, 0, 0)",
    paper_bgcolor="rgba(0, 0, 0, 0)",
    coloraxis_showscale=False
)

# fig.show()

fig.write_image("./metaminer_bioarchiv/abau_comparison/sequencing_technology_normalized.png", format="png", scale=4)
```

In [28]:

```
# match_df = pd.DataFrame({
#     "Category": ["Country", "State"],
#     "Matched": [2496, 2489],
#     "Total": [2500, 2500], 
# })

# # Create bar chart with background (Total)
# fig = px.bar(match_df, x="Category", y="Total", color_discrete_sequence=['#FFFFFF'], barmode="overlay")

# fig.update_traces(marker_line_color="#cccccc", marker_line_width=2)
# fig.update_traces(marker=dict(color="#FFFFFF", opacity=0.7))

# # # Overlay actual progress (Matched)
# fig.add_trace(px.bar(
#     match_df, x="Category", y="Matched", color_discrete_sequence=['']
# ).data[0])

# # Customize layout
# fig.update_layout(
#     title="Geographical Location",
#     xaxis=dict(
#         title="Category",
#         titlefont=dict(size=16, color='black'),
#         tickfont=dict(size=14, color='black'),
#         showline=True,
#         linecolor='black',
#         ticks='outside',
#         tickwidth=2,
#         ticklen=6,
#     ),
#     yaxis=dict(
#         title="# of Assemblies",
#         range=[2400, 2500],  # Lock range to total count
#         tickfont=dict(size=14, color='black'),
#         showline=True,
#         linecolor='black',
#         ticks='outside',
#         tickwidth=2,
#         ticklen=6,
#     ),
#     showlegend=False,
#     height=450,
#     width=500,
#     plot_bgcolor="rgba(0, 0, 0, 0)",
#     paper_bgcolor="rgba(0, 0, 0, 0)",
# )

# fig.update_traces(textposition="inside")

# # Save as PNG
# # fig.write_image("./metaminer_bioarchiv/abau_comparison/geo_loc_normalized.png", format="png", scale=4)

# fig.show()
```
